# Predictive Prioritization of Enhancers Associated with Pancreas Disease Risk

**DOI:** 10.1101/2024.09.07.611794

**Authors:** Li Wang, Songjoon Baek, Gauri Prasad, John Wildenthal, Konnie Guo, David Sturgill, Thucnhi Truongvo, Erin Char, Gianluca Pegoraro, Katherine McKinnon, The Pancreatic Cancer Cohort Consortium, The Pancreatic Cancer Case-Control Consortium, Jason W. Hoskins, Laufey T. Amundadottir, H. Efsun Arda

## Abstract

Genetic and epigenetic variations in regulatory enhancer elements increase susceptibility to a range of pathologies. Despite recent advances, linking enhancer elements to target genes and predicting transcriptional outcomes of enhancer dysfunction remain significant challenges. Using 3D chromatin conformation assays, we generated an extensive enhancer interaction dataset for the human pancreas, encompassing more than 20 donors and five major cell types, including both exocrine and endocrine compartments. We employed a network approach to parse chromatin interactions into enhancer-promoter tree models, facilitating a quantitative, genome-wide analysis of enhancer connectivity. With these tree models, we developed a machine learning algorithm to estimate the impact of enhancer perturbations on cell type-specific gene expression in the human pancreas. Orthogonal to our computational approach, we perturbed enhancer function in primary human pancreas cells using CRISPR interference and quantified the effects at the single-cell level through RNA FISH coupled with high-throughput imaging. Our enhancer tree models enabled the annotation of common germline risk variants associated with pancreas diseases, linking them to putative target genes in specific cell types. For pancreatic ductal adenocarcinoma, we found a stronger enrichment of disease susceptibility variants within acinar cell regulatory elements, despite ductal cells historically being assumed as the primary cell-of-origin. Our integrative approach—combining cell type-specific enhancer-promoter interaction mapping, computational models, and single-cell enhancer perturbation assays—produced a robust resource for studying the genetic basis of pancreas disorders.

## INTRODUCTION

Pancreatic disorders, including diabetes mellitus, pancreatitis, and pancreas cancer, impact over 10% of the global population, placing significant burden on health and economic systems ^1–3^. The pancreas, with its exocrine and endocrine compartments, plays vital roles in both digestion and glucose metabolism. These compartments arise from a common multipotent progenitor during embryonic development and are comprised of distinct cell types, like α-, β-, γ-, duct, and acinar cells ^4,5^. Despite their specialized roles, pancreas cells exhibit remarkable plasticity, with the potential for transdifferentiation and dedifferentiation ^6–8^. While this phenotypic plasticity offers a regenerative potential for replacing lost or injured tissue, it also presents a vulnerability for developing malignancies ^9,10^. For instance, in the case of pancreatic ductal adenocarcinoma (PDAC), growing evidence suggests that acinar cells also contribute to premalignant lesions by transdifferentiating into duct-like states through acinar-to-ductal metaplasia ^11–14^, challenging the long-standing assumption that PDAC originates solely from ductal cells. Therefore, it is crucial to understand the underlying mechanisms that establish and maintain pancreas cell identities ^15–18^, as these mechanisms are likely relevant for regenerative and oncogenic processes.

Enhancers are noncoding DNA elements that regulate gene expression through chromatin interactions and are key regulators in the establishment and maintenance of cell identities. Together with transcription factors, enhancers have an indispensable role in orchestrating tissue-specific gene expression patterns during development, homeostasis, and disease states ^19–21^. Importantly, over 90% of SNPs at disease associated risk loci identified through genome wide association studies (GWAS) are noncoding, with more than 80% of these found in enhancer regions ^22^. However, identifying enhancers, assigning them to their target genes, and determining the specific conditions or cell types in which genetic variants impact enhancer function all remain significant challenges ^23^. This is further complicated by the fact that many enhancers do not activate the closest promoters, and some are located at large distances from their targets ^24^. In the human pancreas, several studies cataloged candidate enhancer regions through open chromatin analysis and epigenetic marks ^25–33^. A few recent studies have initiated efforts to link enhancers to target genes by profiling 3D chromatin interactions in human pancreas cells, however, either their scope was limited to analyzing whole islets without cell-type resolution ^34–36^, or they were constrained by their small sample size ^37,38^.

To address these gaps, we generated cell-type specific, enhancer-promoter interaction datasets using donor pancreas from a comprehensive cohort, spanning 27 donors and five cell types. Overcoming the limitations of the standard pairwise loop analysis, we employed a network approach to parse complex chromatin interactions into tree models, revealing connectivity patterns between enhancers and promoters critical for gene expression. The tree models enabled the development of a machine learning algorithm designed to predict the impact of enhancer perturbations on cell type-specific gene expression, assigning an ‘effect size’ to each enhancer. To validate our predictions and tackle the challenge of measuring cell type-specific enhancer perturbation effects in solid organs like the pancreas, we adapted a high-throughput imaging-based approach to quantify the outcome of enhancer perturbations in single cells from donor tissue. Thus, our study presents a resource for identifying and validating critical enhancers involved in cell type-specific gene expression, while offering a framework for interpreting the genetic basis of complex diseases.

## RESULTS

### Mapping cell type-specific enhancer-promoter interactions using donor pancreas tissue

Enhancer activity is highly cell-type specific. To obtain pure α-, β-, γ-, acinar and duct cell populations from donor pancreas, we refined previously published cell sorting methods, achieving over 95% cell purity in all populations ^29,39,40^ (see Methods, Figure 1A, Supplementary Figure 1A-B). Our protocol was tailored to be compatible with both ATAC-seq and HiChIP assays, facilitating simultaneous chromatin accessibility and 3D chromatin interaction profiling from the same batch of purified cells. Comparing the abundance of key marker gene transcripts in sorted cell populations demonstrated the effectiveness of our cell isolation strategy (Supplementary Figure 1A-B). On these purified cell types, we performed ATAC-seq, and HiChIP using an H3K27ac antibody— a histone modification that marks putative enhancer and promoter elements ^41–45^. These experiments yielded an extensive dataset of 37 ATAC-seq and 29 HiChIP libraries, with each cell type and assay having at least four biological replicates. HiChIP experiments generated more than 5.5 billion reads to allow profiling chromatin interactions at high resolution. Across cell types, we obtained on average, 116,935 accessible regions and 80,947 loops per donor.

**Figure 1.**
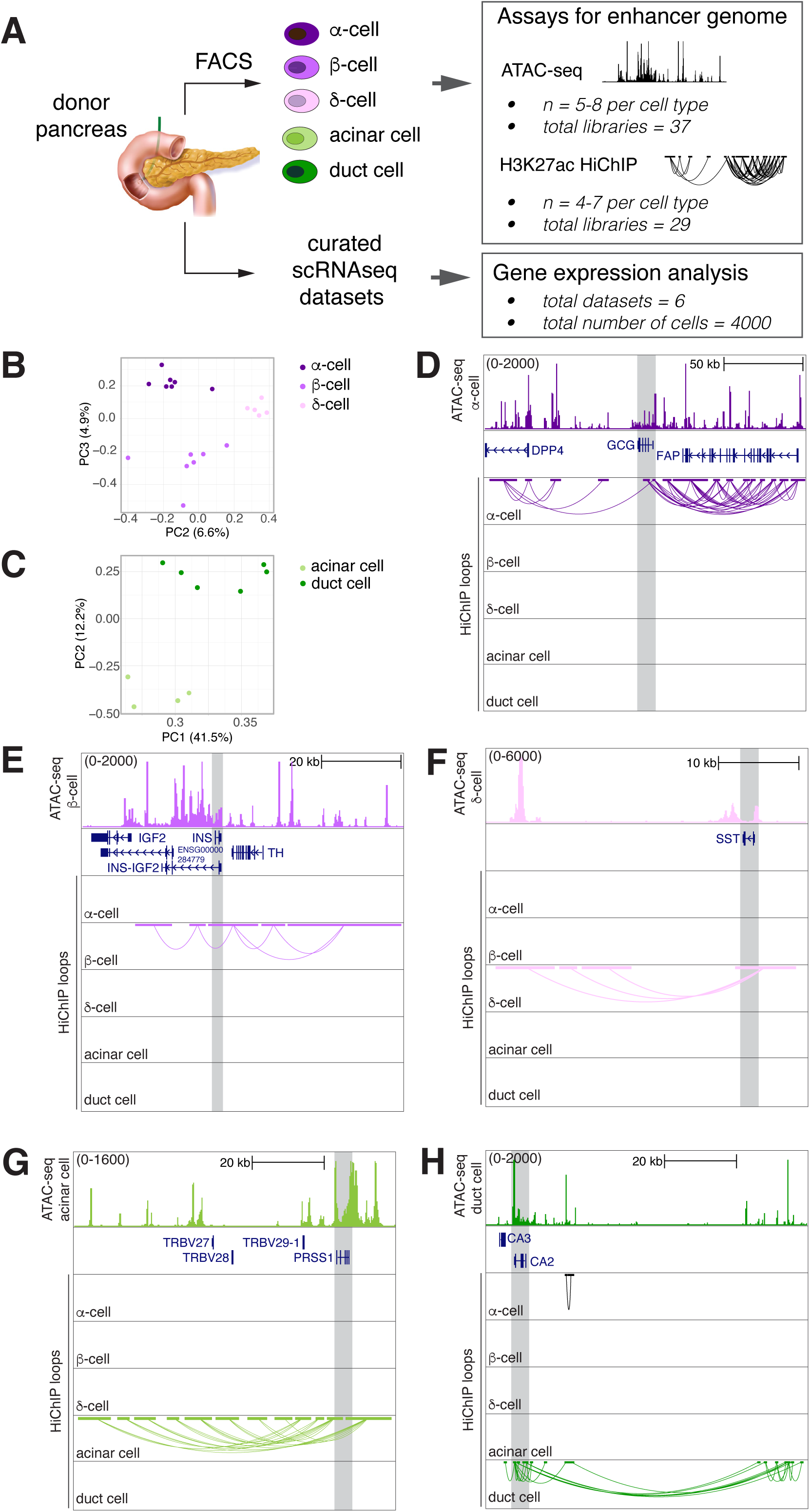
Mapping enhancer-promoter interactions using donor pancreas. A. Overview of the experimental approach. B - C. The PCA plots show the clustering of HiChIP samples based on loop profiles across different pancreatic cell types. Each point represents a sample, and the samples are color-coded according to their respective cell type. D - H. UCSC genome browser tracks showing loci representing cell type-specific HiChIP loops and corresponding ATAC-seq data.

Principal component analysis showed consistent clustering of cell types in our chromatin datasets (Figures 1B-C, Supplementary Figure1C). Looking closely at genomic loci near hallmark genes specific to each lineage revealed loops exclusively associated with the relevant cell types— glucagon (*GCG)* in α-cells, insulin (*INS*) in β-cells, somatostatin (*SST*) in γ-cells, trypsinogen (*PRSS1*) in acinar cells, and carbonic anhydrase (*CA2*) in duct cells (Figure 1D-H). Taken together, our refined cell purification coupled to HiChIP assays captured cell type-specific 3D chromatin interactions in human pancreas cells.

### Parsing enhancer-promoter interactions using graph-based tree models

To gain insight into the 3D chromatin organization in distinct pancreas cell types, we employed a graph-based approach to visualize and analyze HiChIP interactions. In contrast to commonly used visualization methods like chromatin contact matrices or loop arc plots, ‘tree’ graph models facilitate the discovery of hierarchical structures, and the incorporation of other epigenomic data types ^46^.

To build the enhancer-promoter tree models, we first generated a list of consensus loops, representing all loops detected in our combined cell type-specific data (see Methods, Supplementary Figure 2A). We then transformed these chromatin interactions into tree models, where the nodes represent either the enhancer or promoter anchors, and the edges represent chromatin loops detected in our HiChIP experiments. Each tree is defined by its root promoter, therefore can only contain one promoter node (Figure 2A, Supplementary Figure 2B). We began assessing the connectivity with promoters, designating them as the base level— zero (P_0_). Any enhancer directly connected to a promoter was assigned the next tier— level 1 (E_1_). Enhancers that link to E_1_ enhancers, but not directly to the promoters, were then categorized as level 2 (E_2_). Similarly, we designated the loops based on their connectivity (L_1_, L_2_, L_3_, and so forth). This step-by-step assignment continued until all enhancers looping to the promoter were complete ensuring that each enhancer’s level represents its connectivity to the promoter (Supplementary Figure 2B, see Methods for details).

**Figure 2.**
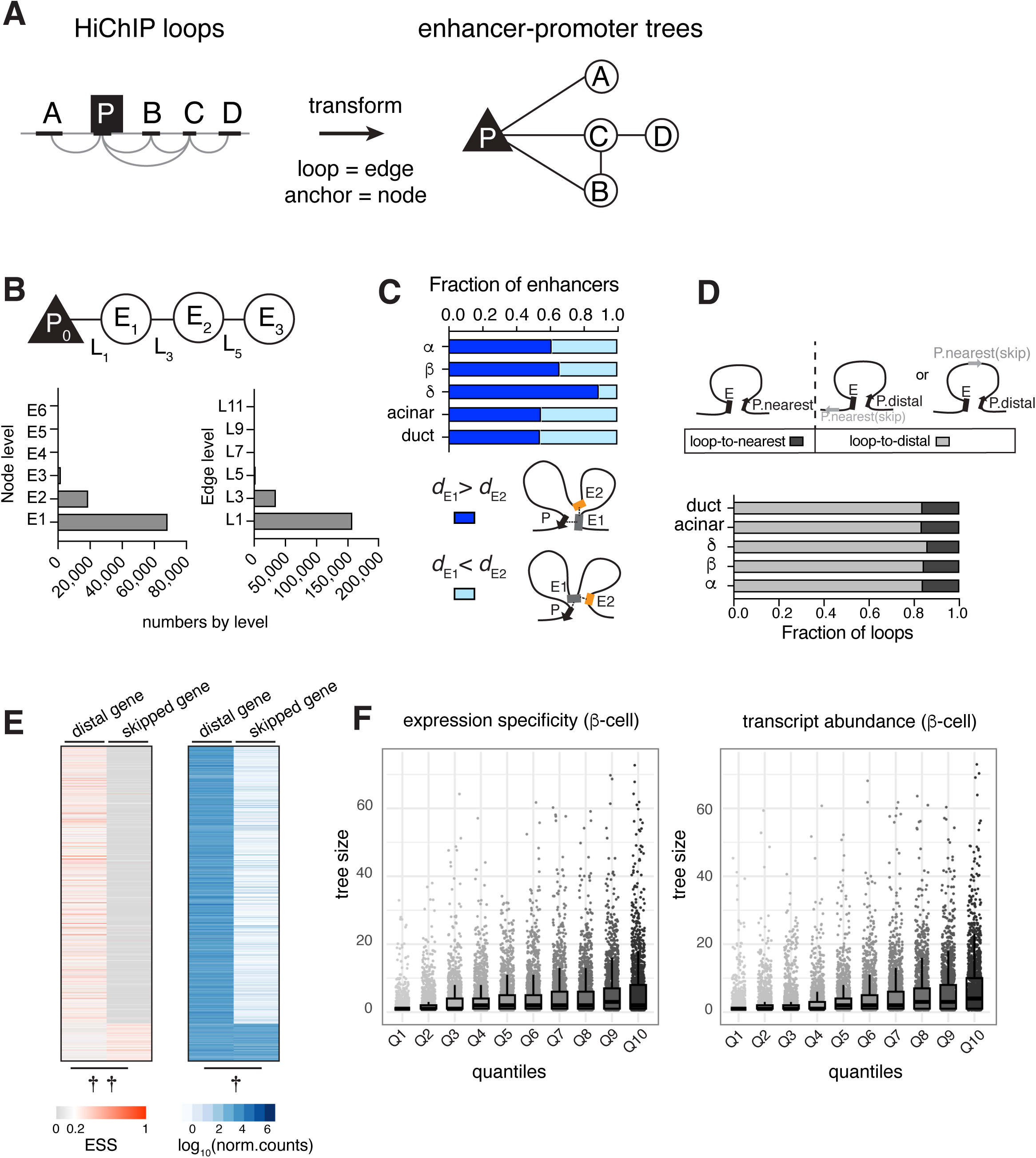
Parsing enhancer-promoter interactions using graph-based tree models. A. Cartoon illustrating how HiChIP interactions are transformed into tree models. Promoter forms the root of the tree, enhancers form the branches, and HiChIP loops are the edges connecting these nodes. B. Enhancers (nodes) and loops (edges) are assigned levels to represent their connectivity within the tree. Bar graphs show the distribution of node and edge numbers by level. Also see Supplementary Figure 2. C. The bar graph shows the proportion of E_1_ enhancers that are located further from the TSS (transcription start site) than E_2_ enhancers (dark blue) and the proportion where E_1_ enhancers are closer to the TSS than E_2_ enhancers (light blue). Possible looping configurations are depicted for each scenario. D. Bar graph shows the fraction of enhancers looping to the nearest gene (dark grey) or a distal gene (light grey) in each cell type. E. Heat maps show the expression specificity (ESS, red) and abundance (blue) of distally looped or skipped genes. Each row represents a gene pair that are either distally looped to or skipped by the same enhancer in β-cells. See Supplementary Figure 2 for graphs of the other cell types. Paired t-test was performed on ESS values or transcript levels of each gene pair, ††*P*-value < 0.0001; †*P*-value < 0.001. F. Box plots depict the relationship between transcript abundance and the size of enhancer-promoter trees as measured by the number of enhancers linked to a single promoter. The x-axis represents the quantiles of expression specificity (ESS) or transcript abundance (normalized counts). The individual data points represent specific tree sizes for genes within each quantile. Data for β-cells is shown. For the plots involving other cell types, see Supplementary Figure 2I.

Parsing the chromatin data into enhancer trees revealed that our HiChIP experiments overwhelmingly captured enhancer-promoter interactions (78%); enhancer interactions that didn’t involve a promoter (orphan enhancers) constituted less than 1% of the data (Supplemental Figure 2B-D). Among the enhancer-promoter interactions, E_1_ enhancers (73%) and L_1_ connections (80%) were the most abundant, suggesting that most enhancers loop to their targets directly (Figure 2B). While categorizing the connections, we also noticed frequent interactions *between* enhancers (i.e. L_2_, L_4_, ∼ 22% of total edges, Supplementary Figure 2D). However, most of these enhancers had a shorter connecting path to a promoter, therefore we further simplified the enhancer trees by pruning these connections (Supplementary Figure 2B). At the end, the indirect loops only constituted ∼11% of all enhancer-promoter interactions.

Since HiChIP assays are based on proximity ligation, we wondered if there is a distance bias for capturing more E_1_ (direct) versus E_2_ (indirect) enhancers. However, when we analyzed the linear distance between these enhancers and their promoters, we found that the E_1_ enhancers were typically located at a greater genomic distance (median 275,552 bp) than E_2_ enhancers (median 149,825 bp, Figure 2C). Further, we assessed the paired-end tag (PET) counts of L_1_ (connecting to E_1_) or L_3_ loops (connecting to E_2_) as a measure for chromatin interaction frequency, and found that L_1_ loops overall exhibited higher PET counts in every cell type, suggesting that E_1_ enhancers likely form more stable loops with their target promoters than other enhancers in the tree, regardless of the distance (Supplementary Figure 2E).

The abundance of E_1_ enhancers prompted us to investigate the functional impact of these direct interactions on target gene expression. We integrated our enhancer trees with gene expression data, compiled from publicly available pancreas single-cell RNA-Seq experiments ^47^. In all five pancreatic cell types, more than 80% of E_1_ enhancers looped to a distal target promoter, bypassing genes closer in linear distance (Figure 2D). We found that the distally looped genes have higher expression specificity and transcript abundance compared to skipped genes (Figure 2E, Supplementary Figure 2F). In addition, more than 68% of skipped genes were annotated as noncoding (Supplementary Figure 2G). This trend persisted even when we limited our analysis to coding genes, with over 60% of E_1_ enhancers still skipping the nearest gene (Supplementary Figure 2H).

We also examined the relationship between enhancer connectivity (tree size) and gene expression. Dividing the expression data into quantiles revealed that genes connected to multiple E_1_ enhancers exhibited higher expression specificity and higher transcript abundance in each cell type (Figure 2F, Supplemental Figure 2I). We speculate that the E_1_ enhancers may collectively promote transcription by increasing the local concentration of lineage-specific transcription factors, leading to robust expression of genes critical for cell identity in each cell type.

### Dissecting enhancer interconnectivity using tree models

In the previous section, we considered the connectivity of individual enhancer-promoter interactions. Enhancers, however, can regulate more than one gene and interact with multiple distinct loci. Standard pairwise loop analysis can underrepresent higher order chromatin contacts or multiway interactions that may exist between multiple enhancer clusters ^48^. Tree models address this by preserving connectivity and dependency between nodes, permitting the discovery of interactions involving more than two regions. Thus, we extended our analysis to include genome-wide enhancer tree interconnectivity in different pancreas cell types.

First, we assessed the extent of enhancers engaged with one or more promoters in our data. Across all cell types analyzed, we observed that on average 60% of enhancers only interacted with a single promoter, 21% interacted with two, and 19% interacted with three or more promoters (Supplementary Figure 3A). Notably, we found that even when an enhancer loops to multiple promoters, its connectivity level rarely changed (Supplementary Figure 3B), suggesting a simpler architecture rather than complex, multi-level enhancer networks. To further explore the enhancer interconnectivity, we focused on enhancer trees that are connected to each other through a shared enhancer or a promoter node. We termed these larger structures ‘enhancer forests’. For this in-depth analysis, we proceeded with α-, β-, acinar and duct cell data, excluding γ-cell data due to the substantially lower number of enhancer trees detected (see Methods).

We found that nearly all enhancer trees belong to a forest— in each cell-type, 92-97% trees are in forests (Supplemental Fig 3C). The median size of a forest includes seven enhancer trees, with a median of 22 nodes, averaged across cell types (Figure 3A). The median genomic span of the enhancer forests is 732kb (Figure 3A), which is similar to the average size of a topologically associated domain (TAD) in the human genome ^49^.

**Figure 3.**
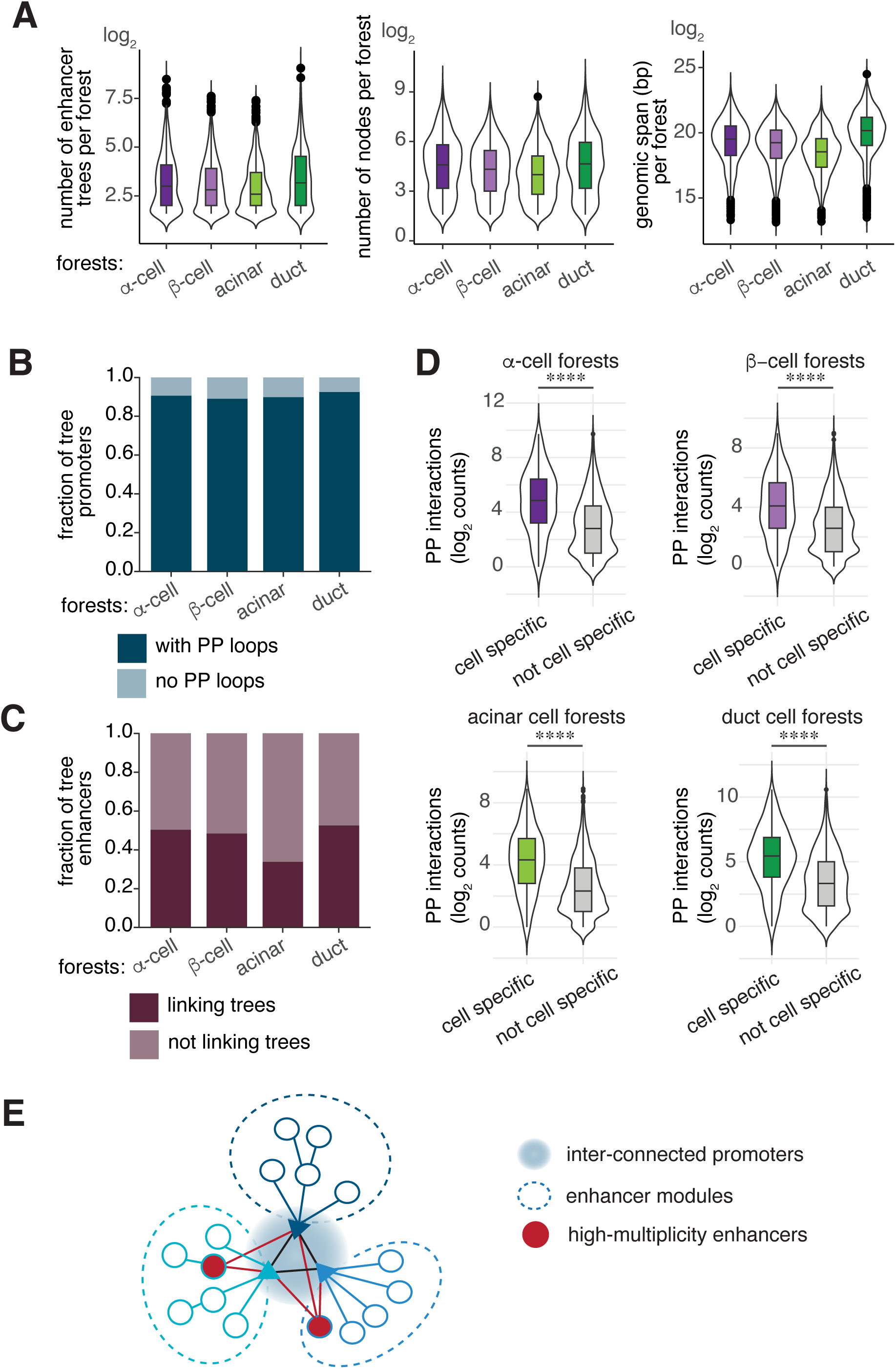
Dissecting enhancer interconnectivity using tree models. A. Violin plots show the distribution of forest features in each cell type. From left to right: number of trees, the number of nodes and the genomic span per forest. B. Bar plots show the fraction of trees whose promoters connect to other trees (with PP loops) and those that do not engage in PP interactions (no PP loops) within each forest. C. Bar plots show fraction of tree enhancers linking to other trees (linking trees) and those that do not (not linking trees) within each forest. D. Violin plots show the distribution of promoter-promoter (PP) interactions in enhancer forests that either contain or do not contain cell type-specific promoters. Mann-Whitney test comparing number of PP interactions in these two categories **** *P*-value <0.0001. E. Model representing the structure of enhancer forests. Promoters that connect different enhancer trees form a central core. Most enhancer nodes within the forest are confined to their own trees forming distinct ‘modules’. Only a few enhancers with high-multiplicity connect multiple trees within the forest, linking these separate modules.

Enhancer trees can form a forest through connections between promoter or enhancer nodes. Analyzing these shared nodes revealed a striking pattern: across all cell types, approximately 90% of promoters connect to another promoter within their forest, whereas only 34%-52% enhancers link distinct enhancer trees (Figure 3B-C). Furthermore, enhancer forests containing cell type-specific genes showed a substantial increase in promoter-promoter interactions with a median of 50 compared to 10 in forests without cell-type specific gene promoters (Figure 3D). The increased connectivity between promoters may facilitate co-regulation of cell type-specific genes to fulfill specialized functions in each cell type. Taken together, these results imply a more central role for promoters in forming the enhancer forests and suggests a modular topology of enhancers where each enhancer cluster typically regulates a specific set of connected genes (Figure 3E).

### Predictive prioritization of cell type-specific enhancers using machine learning

In our investigation of the enhancer-promoter interactions within human pancreas cells, we identified a multitude of enhancers potentially contributing to gene regulation. However, it is unclear if all enhancers contribute equally to gene transcription, or some are more critical than others. We reasoned that our enhancer-promoter trees could facilitate functional prioritization of enhancers, and, importantly, pinpoint those that may underlie disease risk. We developed a machine learning algorithm, named EPIC for *E*nhancer *P*rioritizer using *I*ntegrated *C*hromatin data, capable of predicting the functional impact of enhancers on cell type-specific gene expression. EPIC uses the *k-*nearest neighbor algorithm and our enhancer trees to classify the cell type-specificity of these trees based on chromatin-derived features (Figure 4A). Specifically, we generated a list of 24 predictor variables (six variables per cell type) that include cell type-specific chromatin accessibility (ATAC-seq tags), 3D chromatin interaction frequency (HiChIP PET counts), their interaction terms (ATAC x PET) and the enhancer tree structure (direct vs indirect). EPIC uses these features to learn and make predictions about cell type-specificity of the tree promoters (see Methods for details, Figure 4A). The ground truth class labels for cell type-specificity of these tree promoters were derived from gene expression data obtained through single-cell RNA-Seq studies in the human pancreas ^47^. To assess the performance of EPIC, we evaluated additional models that included 1) only promoter accessibility, 2) enhancer association by linear genomic distance to transcriptional start site and 3) tree model with or without indirect features (Figure 4B, graphic illustration). In all cell types examined, the tree models, which include the direct and indirect interactions, outperformed distance associated or promoter-only models with the highest predictive accuracy (Figure 4C, ROC plots). There were only minor differences between the partial and full tree models, which is expected considering that indirect connections constitute only ∼11% of the data (Figure 2B, Supplementary Figure 2C-D). Notably, we observed that the linear model in α-cells (AUC=0.82) performed nearly as well as the full tree model (AUC=0.84), suggesting a distinct organization of α-cell specific genes in the genome (Figure 4C).

**Figure 4.**
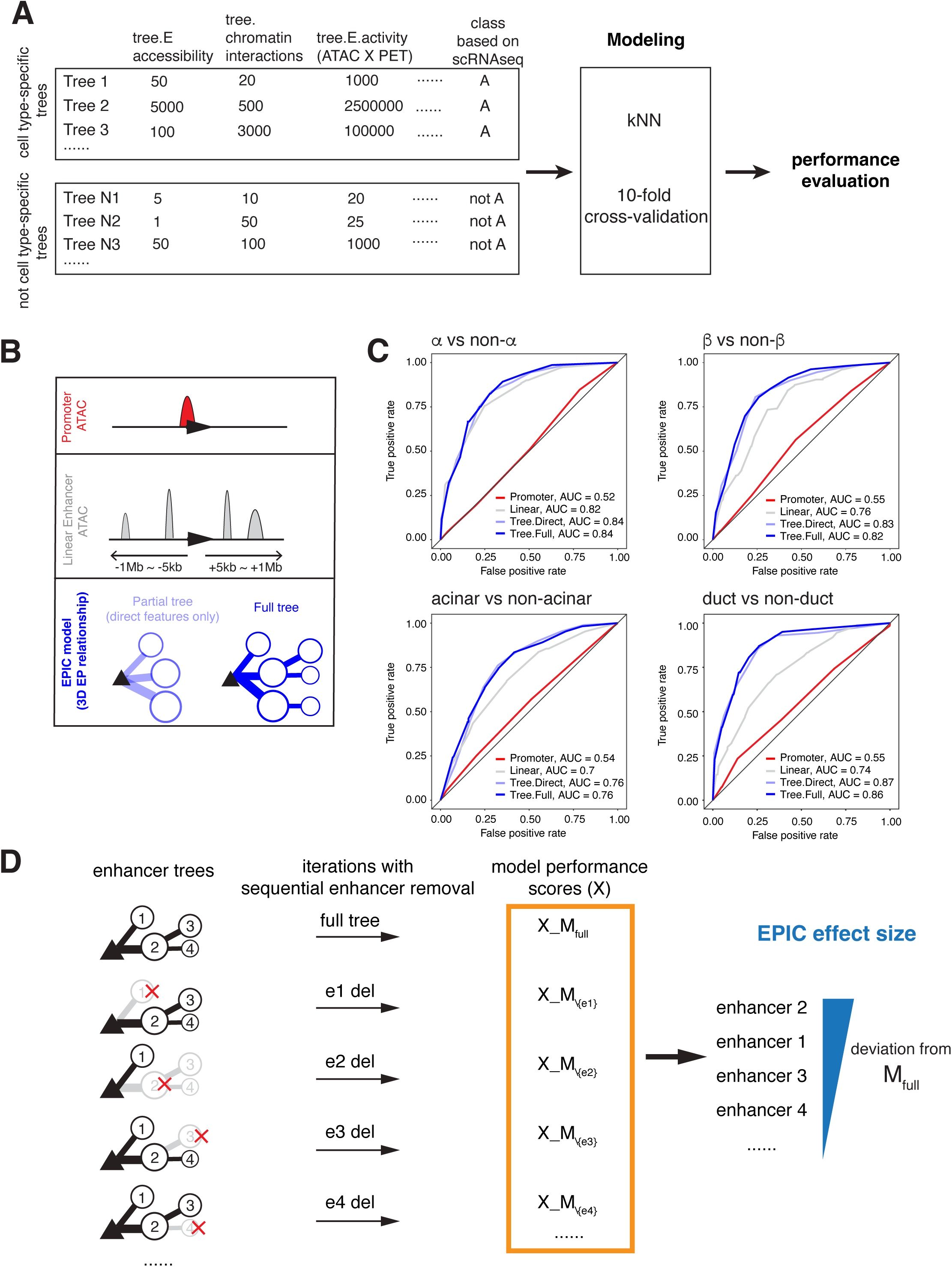
EPIC: A machine learning model to prioritize enhancers impacting cell-type specific gene expression. A. Schematic representation of the data preparation, modeling, and performance evaluation process. Annotated enhancer-promoter trees were divided into two categories: cell type-specific and not cell type-specific, based on expression specificity derived from single-cell RNAseq data. For each tree, chromatin-derived features were calculated, including enhancer accessibility (ATAC-seq), enhancer-promoter interactions (HiChIP PET counts), and enhancer activity (product of accessibility and interaction), and whether the interactions are direct or indirect. The k-Nearest Neighbor (kNN) algorithm was employed to classify trees into their respective cell types using these features. A 10-fold cross-validation was performed to determine the optimal number of neighbors and to evaluate the model’s performance using the accuracy metric. B. Cartoon illustrating alternative models using chromatin features. C. ROC curves showing performance results of models shown in B. The performance of tree-based models was compared to two alternative models: the promoter accessibility model and the linear model. D. Schematic describing *in silico* enhancer perturbations using enhancer trees. Enhancer importance was evaluated by iteratively removing each enhancer from the enhancer-promoter tree and recalculating the model performance (accuracy) using the kNN classifier. The performance of the perturbed model (with a specific enhancer removed) was compared to the full model (with all enhancers). The EPIC effect size for each enhancer was calculated as the deviation in accuracy between the full model and the perturbed model, averaged over multiple iterations. Enhancers with larger deviations were considered to have a greater impact on their associated promoter activity.

Having verified EPIC’s performance, we asked if EPIC could predict the ‘effect size’ of enhancer perturbations for a given target gene. Due to the inherent scalability of graph models, the enhancer trees can flexibly accommodate the addition or removal of nodes (enhancers), and edges (loops). Taking advantage of this feature, we systematically removed each enhancer node and compared accuracy of the enhancer deletion models to the original model in predicting the correct class value— cell type (Figure 4D). Specifically, if an enhancer node was important for cell type-specific expression, its removal would be expected to alter the probability score for that cell type, indicating a reduced confidence in the correct classification and potentially lowering the model’s overall accuracy. Enhancers causing the most significant change in predicted probability were considered to have the highest effect size. This approach allowed us to evaluate, *in silico,* the effect size of perturbations for every enhancer in our enhancer trees.

### Experimentally testing EPIC’s predictions in single cells using donor pancreas

A significant challenge in enhancer perturbation studies using primary human tissue is measuring perturbation effects in a cell type-specific manner, especially in solid organs like the pancreas. To address this, we coupled RNA-FISH with high-throughput imaging ^50,51^ to dCas9-mediated gene activation (CRISPRa) or repression (CRISPRi) ^52,53^ and optimized the assays for donor pancreas cells (Figure 5A, also see Methods). This approach ensured quantitative measurements of mRNAs at the single cell level, in specific cell types, after each enhancer perturbation. Building on our prior success in delivering expression constructs to human pancreas cells ^29^, we introduced adenoviral vectors carrying CRISPRa/i components and guide RNAs (gRNAs) targeting enhancers (see Methods). We typically achieved over 60% transduction efficiency (Figure 5B). After a 5-day recovery period in culture, we performed multiplexed RNA-FISH to quantify the mRNAs of target genes and cell markers (Figure 5A, also see below).

**Figure 5.**
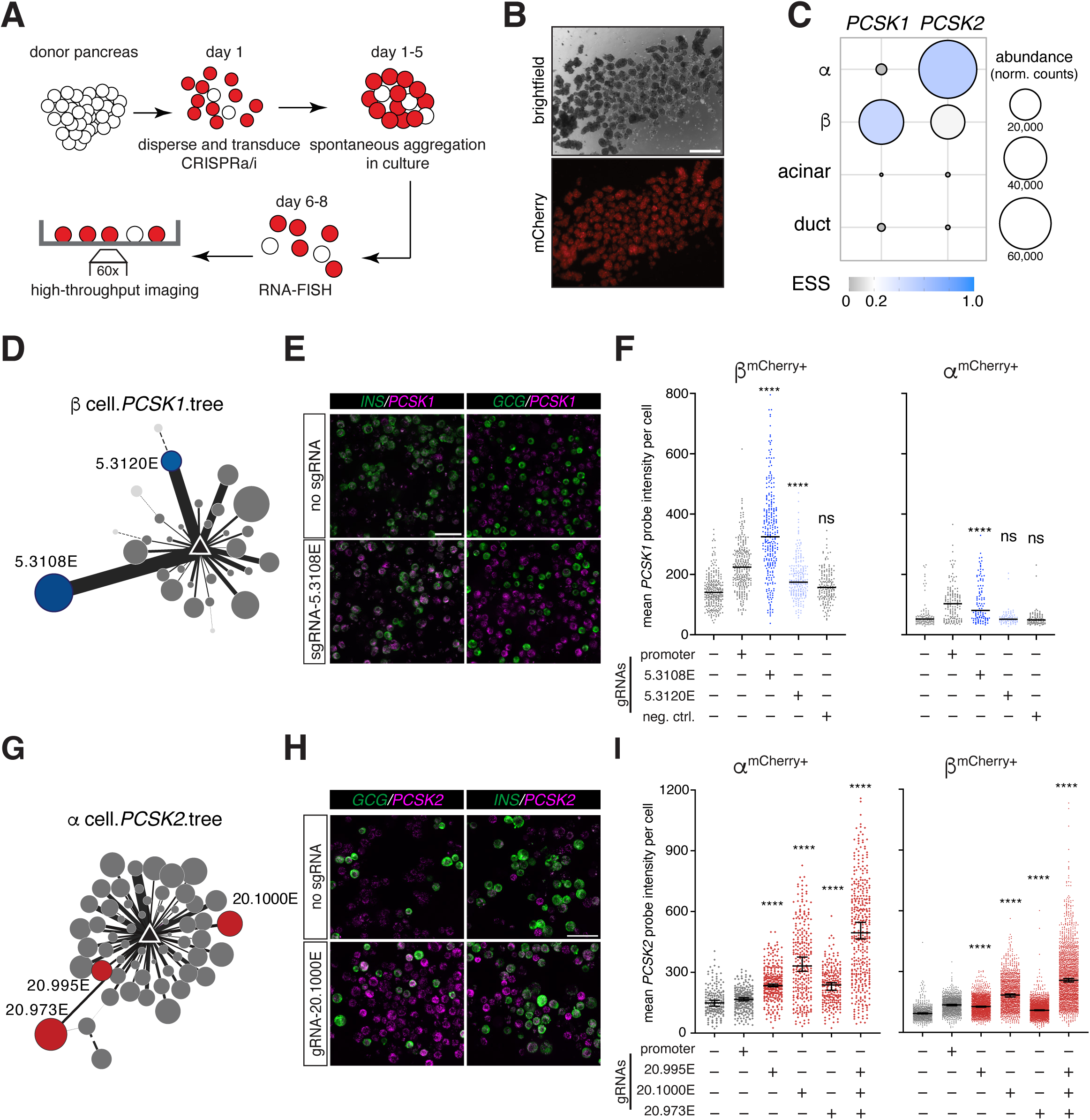
Experimental validation platform of EPIC-prioritized enhancers in single cells using donor pancreas. A. Schematic illustrating the experimental workflow. See methods for details. Red cells represent dCas9-VP64-mCherry^+^ or dCas9-KRAB-mCherry^+^cells after transduction with adenovirus. B. Images showing mCherry^+^ human pancreas cells after transduction. Scale bar: 500μm. C. Bubble plots depict the expression specificity (ESS) and abundance of *PCSK1* and *PCSK2* transcripts in human pancreas cells based on single-cell RNA-seq data. D. Network representation of the β-cell *PCSK1* enhancer tree. Nodes represent enhancers, with the size of each node reflecting the ATAC-seq tag density in β-cells and the thickness of the lines (edges) indicating the strength of the enhancer-promoter interactions as detected by HiChIP. The blue highlighted nodes indicate EPIC-prioritized enhancers. Also see Supplementary Figure 4. E. RNA FISH images showing *PCSK1* transcripts (magenta) in β-cells (*INS* probe, green) and α-cells (*GCG* probe, green) after CRISPRa (dCas9-VP64) modulation. The top row shows cells with no gRNAs, and the bottom row shows cells targeted with gRNAs against the enhancer 5.3108E. Scale bar: 50μm. F. Quantification of *PCSK1* transcript levels in mCherry^+^ β-cells and mCherry^+^ α-cells after CRISPRa (dCas9-VP64) modulation. Mean *PCSK1* probe intensity per cell is shown for cells transduced with gRNAs targeting the promoter (positive control), enhancers 5.3108E and 5.3120E, and a negative control region. The effect of CRISPRa perturbation was analyzed using one-way ANOVA, followed by Dunnett’s multiple comparison test to compare enhancer gRNA targeting conditions to the no gRNA control group. Asterisks indicate *P-*value < 0.0001; ns, not significant. n(mCherry^+^ β-cells)= 1215, n(mCherry^+^ α-cells)= 535, results were reproduced with at least two independent donors. G. Same as D, except the network represents α-cell *PCSK2* enhancer tree, and the red highlighted nodes indicate its EPIC-prioritized enhancers. Also see Supplementary Figure 4. H. Same as E, except *PCSK2* transcript is shown in magenta, and the bottom row shows cells targeted with gRNAs against the enhancer 20.1000E. I. Quantification of *PCSK2* transcript levels in mCherry^+^ α-cells and mCherry^+^ β-cells after CRISPRa (dCas9-VP64) targeting. Mean *PCSK2* probe intensity per cell is shown for cells transduced with gRNAs targeting the promoter, enhancers 20.995E, 20.1000E, and 20.973E. The effect of CRISPRa perturbation was analyzed using one-way ANOVA, followed by Dunnett’s multiple comparison test to compare enhancer gRNA targeting conditions to the no gRNA control group. Asterisks indicate *P-*value < 0.0001. n(mCherry^+^ α-cells)= 1602, n(mCherry^+^ β-cells)= 7477, results were reproduced with at least two independent donors.

To test EPIC’s predictions, we focused on two loci— *PCSK1* and *PCSK2*, due to their hallmark cell type-specific expression (Figure 5C) and their critical roles in hormone processing ^54–58^. Consistent with its role in converting proinsulin to its biologically active form, *PCSK1* expression is most abundant in islet β-cells (Figure 5C). In our HiChIP data, we detected over 30 enhancers forming loops to the *PCSK1* promoter specifically in β-cells, only seven in α-cells, and none in exocrine cells (Supplementary Figure 4A). Because there is no *PCSK1* expression in α- or exocrine cells, we opted to use the CRISPRa system (dCas9-VP64) to examine cell type-specific effects of *PCSK1* enhancer perturbations ^53^. EPIC predicted 5.3108E and 5.3120E as top-ranking prioritized enhancers in β-cells, with 5.3108E having a larger effect size (0.1) than 5.3120E (0.04) (Figure 5D, Supplementary Figure 4B). We designed gRNAs targeting these two enhancer nodes (5.3108E located 32kb, and 5.3120E located 172kb upstream of the transcriptional start site, or TSS). We also designed additional gRNAs targeting the promoter as a positive control and a region 7.6kb upstream of the TSS without any loops as a negative control (Supplementary Figure 4A). After CRISPR targeting, we performed RNA FISH to quantify *PCSK1* transcripts in specific pancreas cells (Figure 5E, *INS* probe marks β-cells, *GCG* probe marks α-cells). As expected, promoter-targeted dCas9-VP64 increased *PCSK1* transcriptional output in β-cells, while the negative control did not significantly alter *PCSK1* levels (Figure 5F). Furthermore, we found that modulating the enhancer regions with CRISPR activation resulted in increased *PCSK1* transcription in the order that EPIC predicted for the effect sizes in β-cells (Figure 5F). Notably, we also observed upregulation of *PCSK1* in α-cells when the top-ranking enhancer, 5.3108E, was targeted (Figure 5F). In contrast, none of the targeted regions activated *PCSK1* in exocrine cells (Supplementary Figure 4C).

*PCSK2* is predominantly expressed in islet α-cells, although single-cell RNA-Seq studies demonstrated *PCSK2* transcript in β-cells ^47^ (Figure 5C), and it has been implicated in the processing of proinsulin to insulin ^59–61^. In line with this evidence, we observed 59 nodes in the *PCSK2* enhancer tree in α-cells, 24 in β-cells, and zero in exocrine cells (Figure 5G, Supplementary Figure 4D). We designed gRNAs targeting three enhancers with effect sizes 0.1 (20.973E), 0.07 (20.1000E), 0.03 (20.995E), and the promoter as a positive control (Supplementary Figure 4B, D). Perturbing these three enhancers with CRISPRa resulted in increased *PCSK2* transcription in α-cells along with the promoter (Figure 5H-I). However, 20.1000E targeting caused the greatest increase even though 20.973E had a slightly higher predicted effect size (Supplementary Figure 4B). Looking into EPIC’s effect size predictions for *PCSK2* enhancers, we noticed that they had a narrow range (0.1 for the highest versus 0.03 for the lowest), potentially revealing regulatory redundancies and consequently the need to perturb more than one element at a time to observe significant alterations to transcription. Indeed, when we targeted all three enhancers simultaneously using CRISPRa, we observed a near-additive effect on *PCSK2* transcript levels (Figure 5I). Perturbing these α-cell specific enhancer tree nodes in β-cells also resulted in upregulation of *PCSK2* transcript, similar to the *PCSK1* findings in α-cells (Figure 5F). These results suggest that the chromatin environment in α- and β-cells, but not in exocrine cells, is permissive to activation at these enhancers. This finding aligns with the fact that both α- and β-cells differentiate from the endocrine lineage during embryonic development.

Taken together, our results demonstrate the potential of our machine learning algorithm, derived from our enhancer tree data, to identify enhancers with a significant impact on target gene transcription.

### Enhancer trees and EPIC facilitate functional annotation of genetic variants associated with pancreas diseases

Hundreds of genetic variants, including single nucleotide polymorphisms (SNPs), have been linked to pancreas diseases by genome-wide association studies (GWAS) ^32,62–64^. However, how these variants impact the disease susceptibility remains largely unclear. Due to linkage disequilibrium (LD), disease-associated loci often contain multiple highly correlated SNPs, making it difficult to identify the causal mechanisms between variants and genes. In addition, determining which cell types are relevant to the disease is challenging since complex diseases, like diabetes or cancer, often involve interactions between multiple cell types. We reasoned that our cell type-specific enhancer-promoter trees in the pancreas, combined with EPIC’s ability to prioritize enhancers, to comprehensively address these challenges and facilitate the functional annotation of genetic variants associated with pancreas disorders.

First, we analyzed the enrichment of GWAS variants related to pancreas disorders within enhancer or promoter nodes across different pancreas cell types using the GARFIELD algorithm ^65^ (See Methods). We found that type 2 diabetes (T2D), and glycemic trait variants were significantly overrepresented in islet cell enhancer trees (T2D in β-cell: *P*=1.52×10^−23^, GWAS threshold 1×10^−07^; fasting glucose in β-cell: *P*=6.99×10^−23^, GWAS threshold 1×10^−05^; HbA1C in β-cell: *P*=1.16×10^−10^, GWAS threshold 1×10^−05^). In contrast, pancreas cancer risk variants were significantly enriched within exocrine cell enhancer trees, with a more significant association observed in acinar cells (*P*=1.47×10^−05^) compared to duct cells (*P*=3.65×10^−03^, GWAS threshold 1×10^−05^) (Figure 6A). As a control, we tested non-pancreas disease related SNPs, like those linked to Alzheimer’s disease or lung cancer. None of these traits showed significant enrichment in the enhancer or promoter nodes in any pancreas cell type (Figure 6A).

**Figure 6.**
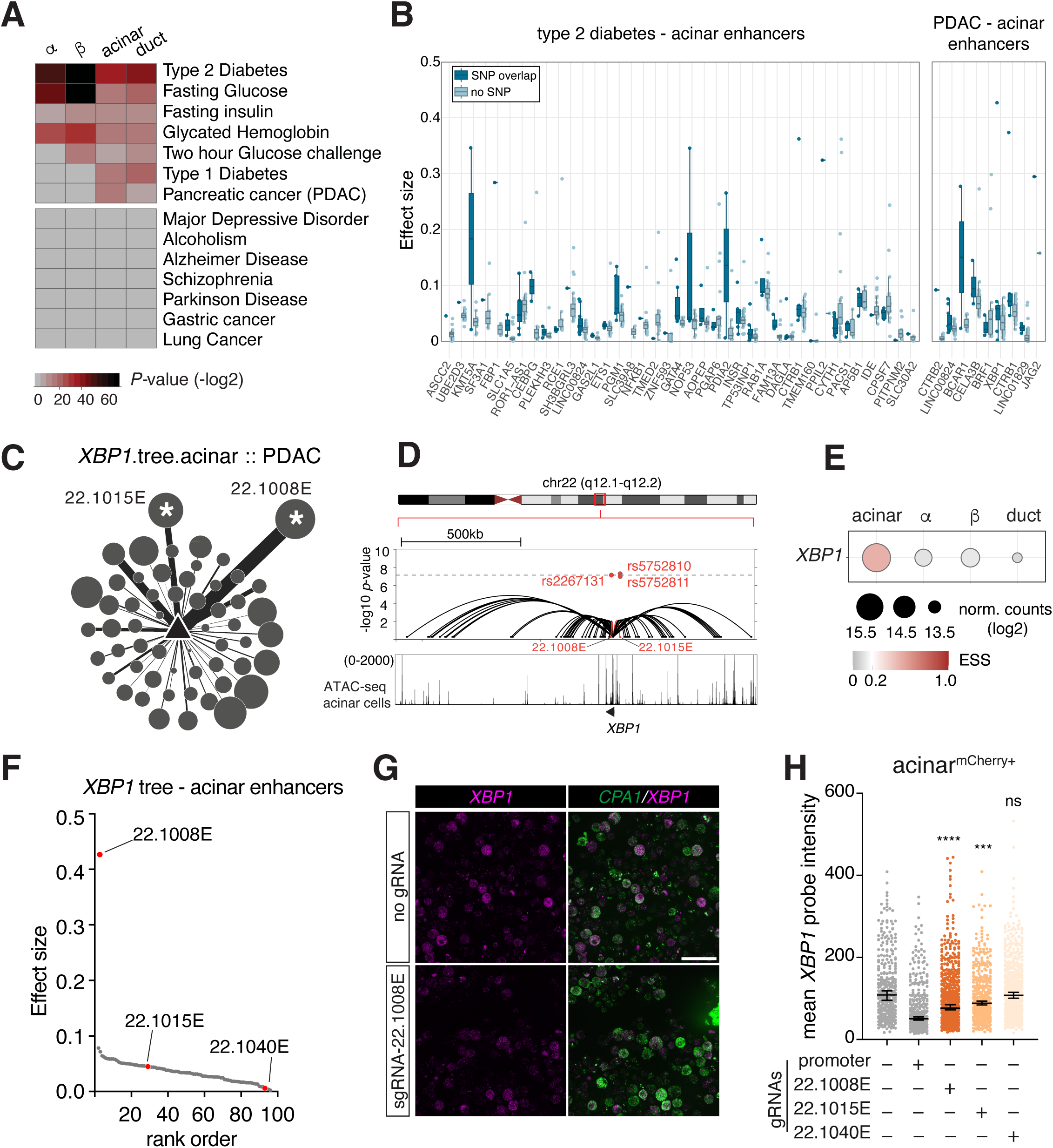
Enhancer-promoter trees and EPIC facilitate annotation of genetic variants linked to disease risk. A. Heat map represents the enrichment analysis results of risk SNPs associated with pancreas disorders and negative control traits in cell type-specific enhancer or promoter nodes. B. Box plots with overlaid data points showing the distribution of effect sizes for acinar cell enhancers associated with type 2 diabetes (left) and pancreatic ductal adenocarcinoma (PDAC) (right). Enhancers are grouped by gene, with dark blue points representing enhancers that overlap with a SNP and light blue points indicating those without an overlapping SNP. The whiskers extend to the most extreme data points not considered outliers. C. Network representation of the *XBP1* enhancer tree in acinar cells. Nodes represent enhancers, with the size of each node reflecting the ATAC-seq tag density and the thickness of the lines (edges) indicating the strength of the enhancer-promoter interactions as detected by HiChIP. The nodes with the asterisks indicate the SNP enriched enhancers. D. Combined UCSC genome browser and locus zoom plots displaying the enhancer tree elements at the *XBP1* locus in acinar cells. The upper panel shows the locus zoom plot with GWAS *P*-values for SNPs associated with PDAC, highlighting significant SNPs (rs2267131, rs5752810, and rs5752811) in red. The UCSC genome browser tracks below show the corresponding ATAC-seq peaks and HiChIP loops detected in acinar cells. Red highlights the nodes enriched with the significant SNPs. E. Bubble plot depicts the expression specificity (ESS) and the abundance of *XBP1* transcripts in human pancreas cells based on single-cell RNA-seq data. *XBP1* is most abundant in acinar cells. F. Scatter plot displaying the ranked effect sizes of acinar cell *XBP1* tree enhancers. The enhancers that were tested in CRISPR perturbation assays are marked in red. G. RNA FISH images showing *XBP1* transcripts (magenta) in acinar cells (*CPA1* probe, green) after CRISPRi (dCas9-KRAB) modulation. The top row shows cells without gRNA treatment, and the bottom row shows cells targeted with gRNAs against the enhancer 22.1008E. Scale bar: 50μm. H. Quantification of *XBP1* transcript levels in mCherry^+^ acinar cells after CRISPRi (dCas9-KRAB) perturbation. Mean *XBP1* probe intensity per cell is shown for cells transduced with gRNAs targeting the promoter (positive control), enhancers 22.1008E (top-ranking), 22.1015E (mid-rank), 22.1040 (low-rank). The effect of CRISPRi perturbation was analyzed using one-way ANOVA, followed by Dunnett’s multiple comparison test to compare enhancer gRNA targeting conditions to the no gRNA control group. ****, *P*-value < 0.0001; ***, *P*-value < 0.001; ns, not significant. n(mCherry^+^ acinar cells)= 2319, results were reproduced by at least two independent donors.

While the enrichment of T2D risk variants in islet cells is expected considering their critical role in glucose metabolism, notably we also observed significant enrichment of T2D risk variants in exocrine cell-specific trees (141 total SNPs, 44 SNPs with GWAS *P*-value < 1×10^−08^, and 68 SNPs with 1×10^−08^ < *P*-value < 1×10^−04^). Closer examination of the tree promoters revealed genes with important functions in acinar cells. For example, germline mutations in the transcription factor *GATA4* have been implicated in childhood onset diabetes and exocrine deficiencies ^66^. Our data placed five T2D risk SNPs in a *GATA4* enhancer at chr8p23.1 (Supplementary Figure 5A-C), linking these variants to a putative gene target in acinar cells.

The enrichment analysis revealed hundreds of SNPs associated with pancreas disorders that overlap with our cell type-specific enhancer or promoter nodes, and our tree models allowed us to link these variants to putative target genes. To test EPIC’s utility in prioritizing enhancers, we compared the effect sizes of enhancers overlapping disease-associated SNPs to those without SNP overlap (T2D and PDAC data in acinar cells shown in Figure 6B). Specifically, for a given cell type and trait association, we identified the SNP-enriched enhancer nodes, retrieved the enhancer trees including those nodes and ranked all the enhancers in the tree based on EPIC’s effect size predictions. We used cumulative density function to determine whether SNP-overlapping enhancers have higher or lower effect sizes compared to enhancers with no SNPs. In the trait-cell type associations we examined, we found that SNP-overlapping enhancers had higher effect sizes than those without SNP overlap (Supplementary Figure 5D-G). This finding suggests that the enhancers prioritized by EPIC may be significant for disease risk.

For pancreatic ductal adenocarcinoma (PDAC), our analysis indicated a stronger enrichment of disease associated variants in acinar cells, even though ductal cells historically have been the focus of PDAC research. We identified three PDAC risk SNPs enriched in two enhancers of *XBP1* at chr22q12.1 in acinar cells (Figure 6C-E). Of these two enhancers, one (22.1008E) overlaps with variant rs2267131 and has the highest EPIC effect size (0.43) for *XBP1* in acinar cells, while the other (22.1015E) has a lower effect size (0.05) and overlaps variants rs5752810 and rs5752811 (Figure 6D, F). We targeted these enhancers using CRISPRi in acinar cells (Figure 6G), including an additional enhancer (22.1040E) that ranked among the lowest as negative control (Figure 6F). We observed a significant reduction in *XBP1* transcripts in acinar cells targeting 22.1008E and 22.1015E, but not in cells targeting the 22.1040E enhancer, demonstrating that the SNP-enriched 22.1008E and 22.1015E enhancers regulate *XBP1* transcription in acinar cells (Figure G-H). Importantly, the degree of reduction was consistent with their EPIC-predicted effect sizes (Figure 6F, H).

Taken together, our chromatin derived enhancer trees help annotate pancreas disease risk SNPs as candidate functional variants influencing high impact enhancers and link them to putative target genes in a cell type-specific manner.

## DISCUSSION

We presented a detailed map and analysis of enhancer-promoter interactions in primary human pancreas cells. By refining cell purification methods and coupling them to ATAC-seq and HiChIP assays, we have generated a high-resolution dataset that captures the enhancer chromatin interactions, surpassing previous work in specificity, sample size and depth.

Our analytical approach is based on versatile, graph-based tree models that simplify the representation and interpretation of chromatin interaction data, permitting systematic quantification of enhancer connectivity. As a result, our analysis demonstrated that direct enhancer (E_1_) interactions are a predominant feature of gene regulation, with multiple enhancers collectively acting on individual promoters to increase transcript output and spatiotemporal specificity. This finding is consistent with previous studies in the developing human cortex ^67^ and mouse embryonic stem cell lineages ^68^, where promoters with significantly high levels of chromatin interactions correlated with lineage specific genes and higher transcriptional levels. However, these prior studies primarily focused on the overall level of chromatin interactions without distinguishing the type of enhancer-promoter connectivity. Our tree models transformed these findings and found that most enhancers that most enhancers form direct loops to their target genes, supporting a simpler, potentially an additive model for enhancing transcription output and specificity ^69,70^. In addition, our forest analysis showed that extensive enhancer sharing between trees is uncommon, with most interconnectivity among trees occurring through promoter-promoter interactions. This suggests that enhancers might be forming distinct modules ^71^ with their respective promoters within the nucleus (Figure 3E), potentially, already positioned for transcription activation when transcription factors reach sufficient local concentrations ^72^.

One caveat of Hi-C assay derivatives like HiChIP is, they capture chromatin interactions as an average across potentially diverse chromatin conformations from millions of cells at a single point in time. This means that the interactions detected may reflect a composite view of different interaction subsets rather than uniform patterns across all cells. Therefore, it remains unclear whether the observed interactions represent persistent enhancer-promoter interactions in every cell, or a sum of varying subsets present in different cells. Emerging single-cell technologies, ^73,74^, may help clarify these distinctions and reveal dynamic enhancer usage based on transcriptional states within the same cell type.

Modifying or perturbing enhancer function is key to understanding enhancer dysregulation in disease ^23^. Prior work on enhancer perturbation primarily used homogeneous cell lines, stem cell-derived cells that have the advantage of unlimited expansion, or whole tissues without cell type resolution ^33,36,75,76^. Here, we used a high-throughput, single cell imaging-based approach to measure the effects of enhancer perturbations in a structurally complex, solid organ like the pancreas.

Our chromatin data combined with tree models form the basis of EPIC, an algorithm designed to prioritize enhancers based on their predicted impact on cell type-specific gene expression. We validated EPIC’s predictions on a select number of enhancer trees and found that our orthogonal imaging-based enhancer perturbation results correspond well with the predicted importance. When analyzing the effect size distributions however, we observed that the differences in EPIC-predicted effect sizes among enhancers within a given tree are relatively small. Furthermore, most enhancer trees contain only a few outlier enhancers whose deletion causes significant deviation of EPIC prediction accuracy, indicating that such critical enhancers are rare. This scarcity of critical enhancers might reflect an evolutionary selection for robust lineage-specific gene expression, where multiple enhancers contribute additively or redundantly to ensure stable regulation. Consistently, our CRISPR-RNA FISH experiments demonstrated this additive effect on *PCSK2* enhancers in α-cells. Further work involving systematic perturbation of combination of elements might help clarify the function of enhancers with smaller effect sizes.

Determining the cell type(s) affected by germline risk variants identified through GWAS, and understanding their specific impact on gene regulation at scale, remain ongoing challenges. Our cell type-specific enhancer trees enabled us to nominate candidate functional variants and target genes at GWAS risk loci across several pancreas diseases. While we confirmed previously known associations, we also identified novel links between risk SNPs and their putative transcriptional targets in understudied cell types ^77,78^. Specifically, we link acinar cells with the overall inherited risk of PDAC. We also observed substantial crosstalk between the exocrine and endocrine compartments of the pancreas in the context of enhancers and risk SNP associations, for instance, the enrichment of type 1 and type 2 diabetes SNPs within exocrine enhancers ^29,79^ This crosstalk suggests a complex interplay between different cell types in disease susceptibility. Future research delineating the contributions of various cell types to these disease etiologies will be crucial for understanding the genetic risk of diseases.

Additionally, our analysis revealed that enhancers overlapping with risk SNPs are more likely to be top-ranking enhancers. Notably, in the trait-cell type associations we analyzed, we observed that some genes have top-ranking enhancers without SNP overlap. We speculate that these enhancers might harbor disease risk variants that are yet to be discovered. In a prior study, HiChIP assays facilitated the identification of promoter-interacting expression quantitative trait loci (pieQTLs) in immune cell types, supporting the view that chromatin features can reveal genetic variants impacting gene expression and disease susceptibility ^80^.

Our study provides new tools and resources for prioritizing enhancers central to cell type-specific gene expression. Exploring the associations we have discovered between genetic variants and their putative gene targets should advance our understanding of how enhancer dysfunction contributes to diseases, and potentially accelerating the development of novel therapeutic strategies.

## METHODS

### Human pancreas tissue procurement

Pancreas tissues were procured through the Integrated Islet Distribution Network (IIDP, USA) and the University of Alberta Islet Core (Canada) from non-diabetic adult organ donors who were deceased due to acute trauma or anoxia. All studies involving human pancreas tissue were conducted in accordance with National Institutes of Health, Institutional Review Board guidelines, and by the Human Research Ethics Board at the University of Alberta (Pro00013094). All donor families provided informed consent for the use of pancreas tissue in research.

### Flow cytometry

#### Preparing human pancreas cells for flow cytometry

Pancreas tissue was shipped to the laboratory by overnight delivery and processed immediately upon receipt without additional culturing time. The tissue was dissociated into single cells following the protocol described in Arda et al 2018 ^29^, with modifications. The tissue samples were pelleted at 200 RCF for 5 minutes and washed once with cold PBS buffer (Thermo Fisher Scientific 10010023) containing 0.1% Pluronic-68 (Gibco 24040-032, PBS-Plu Buffer). The washed tissue pellet was gently resuspended in PBS-Plu Buffer containing 50 μg/mL DNase-I (Sigma-Aldrich DN25-1G) and incubated in a 37°C water bath with gentle agitation for 7-10 minutes. After pelleting, the supernatant was discarded, and the tissue pellet was washed once with cold PBS-Plu Buffer. Next, accutase treatment (Sigma-Aldrich A6964-100ML) was performed to further dissociate the bulk tissue. Islets were resuspended in prewarmed accutase solution at 1000 IEQ/mL concentration, and exocrine tissues at 100 μL (packed pellet)/mL, then incubated in a 37°C water bath for 6-8 minutes with gentle agitation. The accutase was neutralized by adding an equal volume of cold PBS-Plu Buffer. The digested tissues were pelleted and washed once with cold PBS-Plu Buffer. A second round of digestion was performed to achieve single-cell dissociation. The pellets were resuspended in prewarmed PBS-Plu Buffer containing 50 μg/mL DNase I and Dispase (Sigma-Alrich 04942086001) (0.1 U/1000 IEQ for islets, or 1 U/mL for exocrine tissue), and incubated in a 37°C water bath for 6-8 minutes with gentle agitation. The enzymes were neutralized by adding an equal volume of cold FACS Buffer (PBS containing 2% FBS [HyClone, Cytiva], 50 mM EGTA pH 8.0), followed by an additional wash with cold FACS Buffer. Finally, the dissociated cells were passed through a 70 μm mesh (Fisher Scientific 08-771-2) to remove cell debris.

#### Flow assisted cell sorting (FACS)

Dissociated single pancreas cells were resuspended in FACS buffer. Prior to staining with specific antibodies, the cells were incubated on ice for 20 minutes with rat IgG (Thomas Scientific C840F01) (1μL per million cells) to block non-specific binding, and eFluor450 fixable viability dye (Thermo Fisher Scientific 65-0863-18) to label dead cells. Cells were then washed once with FACS buffer and twice with PBS-Plu buffer. Specific antibody staining procedures for exocrine cells and islet cells are as follows:

##### Exocrine cells

We used HPi2-Dylight 650 (Novus, NBP1-18946C) to label and exclude islet cells, HPx1-Dylight 488 (Novus, NBP1-18951G) to label acinar cells and CD133/1(AC141)-PE (Miltenyi Biotec 130-080-801), CD133/2(293C3)-PE (Miltenyi Biotec 130-090-853) to label the duct cells. See Supplementary Figure 1A for the gating strategy.

##### Islet cells

We modified published intracellular staining protocols to make the fixation conditions compatible with downstream ATAC-seq and HiChIP experiments ^40^. Prior to staining with specific antibodies, the cells were fixed in 1% formaldehyde-PBS/Plu buffer (Thermo Fisher Scientific 28906) at 1million cell/mL for 10 minutes at room temperature with rotation. Fixation was stopped by adding glycine solution (125 mM final concentration) for 5 minutes at room temperature. Fixed cells were permeabilized with 1x Permeabilization Buffer diluted in PBS (Thermo Fisher Scientific, 08-8333-56). After permeabilization, cells were stained with INS (D3E7)-biotin (Abcam ab20756) and Streptavidin-APC (Thermo Fisher Scientific 17-4317-82) to label β-cells, GCG(U16-850)-PE(BD565860) to label α-cells, SST-Alexa Fluor 488 (BD566032) to label γ-cells.

All antibodies were used at 1:100 (v/v) concentration, except for HPx1 (2 μL per million cell), INS (1:50) and GCG (1:50) in FACS buffer. All antibody incubation steps were performed on ice for 30 minutes. Labeled cells were sorted on an BD FACS Aria III (BD Biosciences) using a 100 μm nozzle and FACS Diva 8 software, with appropriate area scaling and doublet removal. Gates were determined using fluorescence-minus one (FMO) controls. Sorted populations were collected into low retention tubes containing 100-300 μL cold sort buffer containing 5% BSA (Sigma-Aldrich 126609) in PBS. Cytometry data were analyzed and plotted using FlowJo v.10 (BD Life Sciences).

### Cell purity verification by quantitative RT-PCR (qPCR)

Total RNA was extracted from approximately 5,000 FACS purified cells using the Zymo Direct-zol RNA MiniPrep Kit (R2051) following manufacturer’s instructions. Entire total RNA was used to synthesize the cDNA using the Invitrogen SuperScript III Reverse Transcriptase kit (Thermo Fisher Scientific 18080-044). qPCR reactions were set up using TaqMan gene expression assays and SYBR Green PCR Master Mix (Thermo Fisher Scientific 4368708). The following TaqMan probes were used: *INS* (Hs00355773_m1), *GCG* (Hs00174967_m1), *SST* (Hs00356144_m1), *CPA1* (Hs00156992_m1), *ACTB* (Hs01060665_g1), *CHGA* (Hs00154441_m1), *KRT19* (Hs00761767_s1). Each sample was run in triplicate on a QuantStudio 5 Real-Time PCR machine (Thermo Fisher Scientific). Enrichment of cell marker mRNAs was calculated using the delta(deltaCt) method relative to the mRNA levels in presort cells using the QuantStudio Design and Analysis Software (v1.5.2, Applied Biosystems). To estimate the purity of sorted populations, we used the method described in ^27^.

### ATAC-seq assays

We followed the Omni-ATAC-seq protocol described in ^81^. 5,000-20,000 sorted cells were used for each assay. To isolate the nuclei, cells were resuspended in cold ATAC-Resuspension Buffer (RSB) containing 0.1% NP40 (Sigma-Aldrich I8896), 0.1% Tween-20 (Sigma-Aldrich P1379), and 0.01% Digitonin (Sigma-Aldrich 300410), and incubated on ice for 3 minutes. The lysis was washed out by addition of 1mL of RSB containing 0.1% Tween-20 and mixed by inverting tubes. The nuclei are then pelleted at 500RCF for 10 minutes at 4°C. For transposition, each nuclei pellet was resuspended in 50 μL of transposition mix (25 μL 2x TD buffer, 2.5 μL transposase (100 nM final), 16.5 μL PBS, 0.5 μL 1% digitonin, 0.5 μL 10% Tween-20, 5 μL H2O) and incubated at 37°C for 30 minutes in a thermomixer with 1000 rpm shaking. After transposition, reactions were stopped by adding EDTA (Thermo Fisher Scientific 15575020) to a final concentration of 40 mM. For endocrine samples, reverse crosslink buffer (50 mM Tris, 1 mM EDTA, 1% SDS, 0.2 M NaCl) containing 0.2 mg/mL Proteinase K (Thermo Fisher Scientific 25530049) (add fresh) was added in 5x volume to each reaction and incubated at 65C overnight. Transposed DNA fragments were purified using Zymo DNA Clean and Concentrator-5 Kit (Zymo D4014) and pre-amplified for 5 cycles, for which each reaction contained 2.5 μL of 25 μM i5 primer, 2.5 μL of 25 μM i7 primer, 25 μL 2x NEBNext master mix (NEB M0541), and 20 μL transposed/cleaned-up DNA. PCR conditions (72°C for 5 min, 98°C for 30 sec, followed by 5 cycles of [ 98°C for 10 sec, 63°C for 30 sec, 72°C for 1 min] then hold at 4°C. To determine additional cycles for each sample, 5 μL of the pre-amplified mixture was used to run qPCR in 15 μL reaction containing 3.76 μL sterile water, 0.5 μL 25 μM i5 primer, 0.5 μL 25 μM i7 primer, 0.24 μL 25x SYBR Gold (Thermo Fisher Scientific S11494 in DMSO), 5 μL 2x NEBNext master mix, with the following cycling conditions, 98°C for 30 sec, followed by 20 cycles of [98°C for 10 sec, 63°C for 30 sec, 72°C for 1 min]. Additional cycles were calculated following method by Buenrostro et al 2015 ^82^. The remaining 45 μL of pre-amplified samples were then further amplified accordingly. The PCR condition was 98°C for 30 sec, followed by n cycles of [98°C for 10 sec, 63°C for 30 sec, 72°C for 1 min] then hold at 4°C. Final PCR reactions were purified using the Zymo kit. Libraries were quantified on Aligent TapeStation 4000 and mixed in 1:1 molar ratio, then sequenced on Illumina NextSeq550 to obtain 75bp pair-ended reads.

### ATAC-seq data analysis

We used PEPATAC version 0.8.3 ^83^ with default parameters to process the ATAC-seq FASTQ files. Specifically, reads were aligned to hg38 using Bowtie with the ‘-X 2000’, mitochondrial and blacklisted regions were removed, peaks were called using MACS2 with ‘-f BED -q 0.01 --shift 0 --nomodel’. All samples included in this study surpassed the current ENCODE quality standards for ATACseq data, with TSS enrichment scores of 15-26, and deduplicated aligned reads greater than 14 million per sample. The PEPATAC output bigwig files were used to visualize peaks in the UCSC genome browser.

To generate a consensus peak file, the peak coordinates from the narrowPeaks output files were merged using BEDtools ^84^ with a merge gap of 0 bp. We used the annotatePeaks.pl tool in the HOMER suite ^85^ to generate a count matrix corresponding the peak regions. The peaks were then filtered by excluding those with less than the median ATAC-seq signal for each sample. The final matrix contained a total of 371,234 peaks from 37 ATAC-seq samples. To identify cell type-specific accessible chromatin regions, we used DESeq package ^86^ and performed different cell type or group comparisons: compare between acinar and duct; compare between alpha, beta, and delta; compare endocrine cells (alpha, beta, delta) and exocrine cells (acinar, duct). Peaks that passed the significance threshold of FDR ≤ 1E-06 were selected, resulting in a total of 108,042 differentially accessible peaks. We assigned the cell type-specific clusters to these peaks using k-means clustering with correlation as the similarity metric.

### H3K27ac HiChIP assays

To perform the HiChIP assays, we obtained on average 600,000 purified acinar, duct, β- or α-cells, and 80,000 γ-cells and due to their low abundance. HiChIP libraries were prepared following procedures described in ^87^ with modifications applicable for low cell input material.

Briefly, if the cells were not already fixed prior to sorting, they were fixed in 1% formaldehyde at 1×10^6^ cell/mL concentration for 10 minutes at room temperature with rotation, quenched with glycine at final concentration of 125 mM for 5 minutes, then pelleted at 500RCF for 5 minutes and washed once with PBS/Plu buffer.

#### in situ contact generation

Fixed cells were resuspended in 250 μL of ice-cold Hi-C Lysis Buffer and rotate at 4°C for 30 minutes. The nuclei were pelleted at 2500RCF for 5 minutes and washed with 250 μL of ice-cold Hi-C Lysis Buffer. The washed nuclei pellet was resuspended in 50 μL of 0.5% SDS (Thermo Fisher Scientific 15553027) and incubated at 62°C for 10 minutes, then quenched by adding 146 μL of H2O and 25 μL of 10% TritonX-100 (Sigma-Aldrich T8787) with incubation at 37°C for 15 minutes, rotating end-to-end. To digest the chromatin *in situ*, 100U MboI (Sigma-Aldrich R0147) and 25 μL 10xNEB buffer 2 (NEB B7002 RT) were added to the reaction and incubated at 37°C for 2 hours with rotation. Enzymes were heat-inactivated at 62°C for 20 minutes. To label ends of the digested chromatin fragments, 14 μL of dNTP (NEB N0446S) (dATP-14-biotin and dTTP, dCTP, dGTP at a final concentration of 40 μM each) and 15U of DNA Polymerase I, Large (Klenow) Fragment (NEB, M0210) were added and incubated at 37°C for 45 minutes. Ends were ligated by adding 484 μL of ligation mixture containing 75 μL 10x T4 ligase buffer, 62.5 μL 10% TritonX-100, 3.75 μL 20 mg/mL BSA, 2000U of T4 ligase (NEB, M0202S) and 337.75 μL H2O and incubated at RT for 2 hours with rotation. The ligation mix was pelleted at 2500RCF for 5 minutes at 4°C.

#### H3K27ac-ChIP

The *in situ* Hi-C pellet was lysed in 130 μL of Nuclei Lysis Buffer then transferred to an AFA microtube (Covaris 520045) and sonicated on a Covaris E220 under the following setting: PIP 105W, duty factor 2%, CPB 200, time 4 minutes, temperature 6°C. The sheared lysate was cleared by centrifugation at 16100RCF, 4°C for 15 minutes. Cleared lysate was diluted 2x in ChIP Dilution Buffer and precleared with 15 μL Dynabeads Protein A (Thermo Fisher Scientific 10001D) at 4°C for 1 hour with rotation. 1 μL of anti-H3K27ac (Abcam ab4729 Lot# GR3211959-1) was added to the precleared lysate to immuno-precipitate (IP) H3K27ac associated, proximity ligated chromatin fragments overnight at 4°C with rotation. 15 μL Dynabeads Protein A were added and incubated at 4°C for 2 hours to pull down the IP-ed complex. Beads were washed in order of the following: Low Salt, High Salt and LiCl (Sigma-Aldrich L7026) buffers. Washing was performed at room temperature on a magnet stand by adding to a sample tube 300 μL of a wash buffer, turning the tube 180 degree relative to the magnet for few times, allowing the beads to set for 2 minutes then removing the supernatant. ChIP DNA was eluted by incubation in 50 μL ChIP Elution Buffer on a thermomixer (Eppendorf) at 37°C with 300 rpm mixing. Two elution cycles were performed per sample and the eluates were pooled. To reverse crosslinks, 5 μL Proteinase K (Thermo Fisher Scientific 25530049) was added and the mixture was incubated at 55°C for 45 minutes followed by 67°C for 2.5 hours. The DNA was purified by the Zymo kit (Zymo D4014) following the manufacturer’s instructions, and eluted with 12 μL water, of which 2 μL was used for quantitation on Aligent TapeStation 4000.

#### Biotin pull-down and sequencing library preparation

5 μL Streptavidin C-1 beads (Thermo Fisher Scientific 65001) were washed with 300 μL Tween Wash Buffer, resuspended in 10 μL 2xBinding Buffer and mixed with each sample. The mixtures were incubated at RT for 15 minutes with rotation. Beads were washed in 300 μL Tween Wash Buffer twice at 55°C with shaking (400 rpm), followed by one wash in 100 μL 2x TD Buffer. For on bead tagmentation, beads were resuspended in 50 μL mixture containing 25 μL 2xTD buffer, 0.05 μL Tn5 (IIlumina 20034198) per 1ng post-ChIP DNA and H2O and incubated at 55°C with interval shaking for 10 minutes. The tagmented beads were incubated in 300 μL 50 mM EDTA at 50°C for 30 minutes, followed by washes in 2x 50 mM EDTA (300 μL) at 50°C for 3 minutes, 3x Tween Wash Buffer (300 μL) at 55°C for 2 minutes then once in 200 μL 10 mM Tris. To prepare sequencing library, beads were resuspended in 50 μL PCR mix (25 μL 2xNEB HF master mix (NEB M054), 1 μL 12.5 μM Nextera ad-noMx, 1 μL barcoded 12.5 μM Nextera ad2.x and 23 μL H2O) and amplified by PCR program: 72°C for 5 min, 98°C for 1min, followed by n cycles of [98°C for 15 sec, 63°C for 30 sec, 72°C for 1 min] then hold at 4°C. N was determined by the amount of post-ChIP DNA as described in Mumbach protocol ^87^. The libraries were cleaned up using the Zymo kit and eluted in 15 - 18 μL H2O, 2 μL of which was used to determine the quantity and size distribution on an Agilent TapeStation 4000. High quality libraries were sequenced with 2×75bp runs on an Illumina NextSeq instrument.

### HiChIP data analysis

Paired-end reads from 29 HiChIP samples were aligned to hg38 using the Hi-C Pro pipeline ^88^ with default settings. Hi-C Pro trims, aligns, assigns reads to MboI restriction fragments, filters for valid interactions and generates binned interaction matrices.

#### Calling loops

We used hichipper ^89^ and FitHiChIP ^90^ to call loops on the Hi-C Pro processed samples. For hichipper, we used the peak calling options *EACH, SELF* which include each sample individually and only self-ligation reads. hichipper interactions were filtered based on a PET count ≥ 2 and a Mango FDR ≤ 0.01, which are classified as “significant loops”. For FitHiChIP, we used the “loose” parameter and a bin size of 5000 kb to call loops at FDR ≤ 0.05. Loops were converted to *bigBed* format then uploaded to UCSC browser for visualization. To evaluate variability across donor samples, PET counts from individual donor loop sets were used as input for the principal component analysis (PCA). To calculate and visualize the PCA of donor samples, the R prcomp() function and the ggplot2 package were used.

#### Generating consensus loops

To construct this loop set, we merged the FASTQ reads of the 29 HiChIP samples based on cell type (α-cell, β-cell, γ-cell, acinar, and duct). We processed the merged reads using the Hi-C pro pipeline, and performed loop calling using hichipper and FitHiChIP as described above. We then intersected the hichipper and FitHiChIP loops for each cell type, using the BEDTools pairtoPair function, which reports the overlaps if two loops have the same anchors. We considered these common loops ‘high confidence loops’ as they were identified by two independent loop callers. To obtain a final list of non-overlapping anchors, we merged 1,144,820 anchors from high-confidence loops across five cell types using BEDTools with the default ‘any overlap’ option, resulting in 127,487 consensus anchors. For each consensus anchor, the midpoint was determined by calculating the average of the start and end positions of the merged anchors, rounded down to the nearest integer. A consensus loop set was also created, in which each unique consensus loop was logged, along with corresponding PETs in every cell type. The length of consensus loop was defined as genomic distance between anchor midpoints. After removing loops with length less than 5 kb (artifact of anchor merging process), we obtained a total of 349,749 consensus loops from all five cell types.

The *AnnotatePeaks.pl* function of Homer suite (version 4.11.1) ^85^ was used with hg38 build (gencodev42 (GRCh38.p13)) to annotate anchors. We defined the promoter regions as 2 kb upstream and 3 kb downstream of the transcription start site (TSS). If the midpoint of an anchor falls into a promoter region, that anchor became a promoter anchor (P), otherwise, an enhancer anchor (E). Enhancer anchors were named using chromosome location, a numerical identifier, and the letter “E” (e.g., 22.1008E). Promoters followed a similar notation, with the letter “P” used instead.

### Constructing the enhancer tree models

Principal network elements were defined using consensus loops. Anchors were represented as nodes and the loops connecting the anchors were represented as network edges.

#### Notations

*N* represents the set of all nodes.

*E* represents the set of all edges (loops).

*L_N_*(*n*) represents the level of node *n*.

*L_E_*(*e*) represents the level of edge *e*, where *e* connects nodes *n*_1_ and *n*_2_.

For an edge *e* connecting nodes *n*_1_ and *n*_2_, let *e* = (*n*_1_, *n*_2_).

#### Assigning levels to nodes

For each iteration until no new nodes are assigned:

For each edge *e* = (*n*_1_, *n*_2_) with either *L_N_* (*n*_1_) ≠ − ∞ or *L_N_*(*n*_1_) ≠ −∞

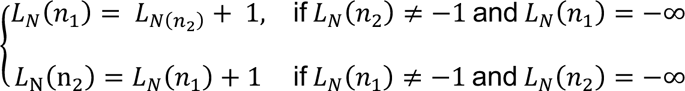

#### Assigning levels to edges

For each edge *e* = (*n*_#_, *n*_$_),

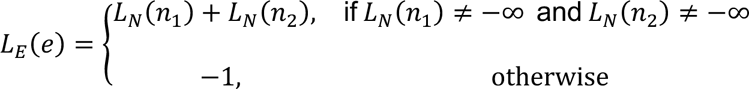

Nodes are assigned levels based on their proximity to a starting node (nodeid). The starting node is assigned a level of 0. In each subsequent iteration, adjacent nodes are assigned a level that’s one greater than the current node. The level of an edge (or loop) is determined by the sum of the levels of the nodes it connects. If either of the nodes has not been assigned a level, the edge remains unassigned (level = -∞).

#### Redundancy Removal

Delete edges *e* = (*n*_1_, *n*_2_) with *L_N_*(*n*_1_) = *L_N_*(*n*_2_).

Consensus enhancer trees were built following these steps described above. Cell type-specific trees were derived by removing the nodes and edges absent from a given cell type. We have obtained 15557 non-trivial enhancer trees for α-cells, 14276 for β-cells, 10949 for acinar cells, and 12483 for duct cells. Forests were identified as two or more enhancer trees connected via shared nodes. To annotate these trees, we integrated the ATAC tag density to the nodes, and the HiChIP PETs to the edges of the cell type-specific enhancer trees.

### Evaluation of enhancer importance by *in silico* perturbation (EPIC)

To estimate the effect size of individual enhancers in an enhancer-promoter tree, we developed a machine learning method to score enhancers by iteratively removing one enhancer at a time and tracking the change in the accuracy of the trained model.

#### Preparing the input data

We selected four sets of enhancer-promoter trees corresponding to four cell types— α-cell, β-cell, acinar, and duct). Data for γ-cell was excluded due to substantially fewer number of enhancer trees detected (Supplemental Figure 2A). γ-cells are among the least abundant cell types within islets (less than 100,000 cells per donor). While we were able to detect frequently occurring loops, like the *SST* locus, the low cell number precludes constructing HiChIP libraries with complexity comparable to the other cell types. We used the expression specificity scores (ESS) ^47^ to subset enhancer-promoter trees that represent both cell type-specific and not cell type-specific genes. Based on our prior work, ESS greater than 0.7 indicates high cell type-specificity, whereas ESS less than 0.3 indicates low specificity ^47^. In the α-cell dataset, there were 354 trees whose associated genes had an ESS greater than 0.7, therefore these trees were assigned the class value “alpha”. To ensure a balanced representation of the non-specific class, we randomly selected a similar number of genes whose ESS is less than 0.3 in all four cell types and assigned them as “non-alpha”. We applied a similar approach for the other cell types. For β-cell versus non-β-cell, we used 340 trees for “beta” and 365 for “non-beta”. The acinar versus non-acinar comparison included 641 trees for “acinar” and 652 for “non-acinar”. Finally, for duct versus non-duct, we used 476 trees for “duct” and 447 for “non-duct”. We named these sets as Salpha, Sbeta, Sacinar, and Sduct. We then built a matrix containing predictor variables based on our chromatin interaction and accessibility data corresponding to these selected enhancer trees. Specifically, for a given tree that belongs to the cell-type A, the sum of ATAC-seq tag densities of direct enhancer nodes (ATAC-d-A), the sum of HiChIP PET counts of direct edges (PET-d-A), the sum of their products (ATACxPET-d-A), and similar 3 variables for indirect nodes (ATAC-i-A, PET-i-A, and ATACxPET-i-A). In total, 24 variables of chromatin derived data were included in the matrix.

#### Initial modeling and performance evaluation

We employed the k-Nearest Neighbor (kNN) algorithm to classify the cell types that the enhancer tree promoters belong based on the predictor variables detailed above. The data was centered and scaled to normalize, and Euclidean distance was used to measure the distance between data points. 10-fold cross-validation was employed to identify the optimal number of neighbors (k). We built the original models Malpha, Mbeta, Macinar, and Mduct on the Salpha, Sbeta, Sacinar, and Sduct sets. To evaluate the model performance, we used the accuracy metric, which is the ratio of correct observations (true positives) to the number of total observations.

#### Comparison of the original tree model performances to alternative genomic models

We compared the performance of tree models against two other models: Promoter accessibility model and the linear model. Promoter accessibility model was built by taking only ATAC-seq tag counts at the promoters and no further chromatin information such as PET counts or interacting enhancers was considered. We defined the promoter regions as 2 kb upstream and 3 kb downstream of the transcription start site (TSS). In the linear model, we assigned a ‘gene regulatory domain’ that extends 1 Mb both upstream and downstream of the TSS. For this model, only the ATAC-seq tag counts of enhancers within this domain were considered, and the ATAC-seq density of each neighboring enhancer within such window was scaled to be inversely proportional to its genomic distance to the promoter.

#### Enhancer removal and evaluating the perturbed model performances

For each enhancer node ‘e’ of the tree ‘T’, we created perturbed models M_alpha_\{e}, M_beta_\{e}, M_acinar_\{e}, and M_duct_\{e} by removing the enhancer node ‘e’ and its child nodes. All 24 predictor variables (ATAC-seq tags, HiChIP PET counts, their products [ATACxPET] for four cell types, direct/indirect node) were recalculated after each enhancer removal. Similar to the original models, we used the kNN algorithm with 10-fold cross-validation splits. By comparing the accuracy difference between the original model and the perturbed one, we calculated the deviation of prediction accuracies between e-deleted tree and the original prediction for each enhancer tree set. By summing up the absolute values of these accuracy deviations across cell types, the total deviation of the enhancer e is given as

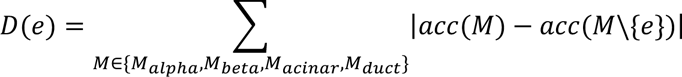

where acc stands for the accuracy of the learning model. We repeated these procedures 100 times for each enhancer to obtain a robust estimate of the average total deviation. This average was defined as the ‘effect size’ for tree T. Enhancers with larger total deviations were considered to have a greater impact on their associated promoter activity. We implemented EPIC algorithm in R, using the caret library for model performance evaluation.

### Perturbing enhancers using CRISPR interference in human pancreas cells

#### CRISPR guide RNA design

CRISPR guides were designed using the software Geneious Prime 2023.0.1(https://www.geneious.com) and were selected based on activity ^91,92^ and specificity scores that exceeded 94% ^93^. We targeted each anchor region (enhancer node) with minimum three gRNAs.

#### Production of adenoviral vectors and viral stocks

Adenoviruses carrying the CRISPR constructs were provided by Vector Biolabs using their custom adenovirus construction service (https://www.vectorbiolabs.com/). Ad-CMV-NLS-dCas9-VP64-mCherry (CRISPRa) was purchased from Vector Biolabs. For CRISPRi, Ad-CMV-NLS-dCas9-KRAB-mCherry were cloned by modifying the SFFV-dCas9-KRAB-BFP (addgene46911) at Vector Biolabs then packaged into adenoviruses. For gRNA viruses, gRNA arrays for each target region were assembled into CARGO constructs following the previously described protocol ^94^, these constructs then were sent to Vector Biolabs to produce adenoviruses. All virus particles were produced at a minimum titer of 1 x 10^10^PFU/ml (PFU, plaque forming unit).

#### Adenoviral transduction of primary human pancreas cells

Human islets or exocrine tissues were partially dissociated using enzymatic digestion as described previously (Accutase, Thermo Fisher Scientific) ^39^. Dispersed cells were seeded on AggreWell400 wells (STEMCELL 34415) at an estimated 300,000-500,000 cells per microwell containing the appropriate culture media. The islet culture media consisted of RPMI 1640 with Glutamine, supplemented with 10% fetal bovine serum (FBS) (HyClone) and 1% penicillin/streptomycin (P/S) (Thermo Fisher Scientific 15140122). The exocrine culture media was composed of CMRL 1066 supplemented with 10% heat-inactivated FBS, 1% GlutaMAX (Thermo Fisher Scientific 35050061), 1% P/S, and 0.1% nicotinamide (Sigma-Aldrich 72340). Cells were transduced with adenovirus at an MOI of 100 and cultured at 37°C for 2 hours. After the incubation, viral media were removed, cells were washed with PBS twice and returned to the incubator with fresh media to allow aggregation. Post-transduction, the cell clusters were transferred to a 24-well ultra-low attachment plate (Corning 3473) on day 3. The media was refreshed as needed, and the cells were harvested for hybridization chain reaction (HCR) preprocessing on day 5.

### RNA-FISH using hybridization chain reaction (HCR) in primary human pancreas cells

CRISPR treated and control cell clusters were collected and dissociated into monolayer using enzymatic dispersion (Accutase, Sigma-Aldrich A6964). After fixation with 4% paraformaldehyde at room temperature for 1 hour, the cells were washed and stored in 1% BSA in PBS with Riboblock (Thermo Fisher Scientific EO0384) at 4°C until further processing. For permeabilization, cells were treated with PBS-Tween20 (PBS-T) containing Riboblock, followed by storage in 70% ethanol at −20°C. The hybridization was performed with probes specific to the targeted genes, followed by a series of washes to remove unbound probes. Amplification of the hybridized probes was carried out using hairpin oligonucleotides, and the samples were stained with DAPI for nuclear visualization. Hybridization experiments were performed using HCR RNA-FISH bundles from Molecular Instruments, including customized probe sets, amplifiers, and buffers specific for single cell suspension and staining. For each sample, multiplexed staining was performed using combinations of target gene probes and marker gene probes. Matched fluorescently labeled amplifier hairpins enabled specific quantification of gene transcripts within the appropriate cell types. The cells were counterstained with DAPI (4′,6-diamidino-2-phenylindole) and seeded at 70–90% density (50,000–75,000 cells per well) on collagen-coated, optically clear 96-well imaging plates (Revvity Health Sciences, D6055700) immediately prior to imaging.

#### High-throughput imaging and quantitative analysis

HCR samples were imaged in four channels (405, 488, 561, and 640 nm) using a dual spinning disk high-throughput confocal microscope (Yokogawa CV7000 or CV8000). A 60x water immersion objective (NA = 1.2) was used for imaging monolayer samples. Two 16-bit sCMOS cameras were employed with binning set to 1, yielding a pixel size of 108 nm. Image Z-stacks were acquired at 0.5-micron intervals across a total depth of 8 microns. For each well, 20–48 randomly selected fields (containing approximately 10–100 cells per field) were imaged. Images were automatically corrected in real-time using Yokogawa’s proprietary software to address camera alignment, optical aberrations, vignetting, and camera background issues. The maximally projected and corrected images were saved as 16-bit TIFF files. Quantitative analysis of the HCR results was done using Columbus 2.9.1 (PerkinElmer/Revvity) or Signal Imaging Artist (SiMA, PerkinElmer/Revvity) 1.2 software. The mean fluorescence intensity in the far-red channel was measured over the cell body region for each cell, and this value was used as the primary output measurement. Single-cell results were exported from Columbus or SiMA as tabular text files. Statistical analysis and data plotting was done in GraphPad Prism version 10.2.3 (GraphPad Software, USA, www.graphpad.com).

### Enrichment analysis of GWAS SNPs risk for pancreas diseases in cell specific EP trees

#### GWAS data selection

GWAS summary statistics for traits associated with pancreas disorders and control traits were obtained from various key studies: Type 2 diabetes ^64^, fasting glucose, fasting insulin, Glycated Hemoglobin and two-hour glucose ^63^ Type 1 diabetes ^32^, and Pancreatic ductal-adenocarcinoma (PDAC) ^62^, control traits ^95,96^. GWAS summary statistics were downloaded from Type 2 diabetes Knowledge portal (https://t2d.hugeamp.org/), MAGIC consortium (http://magicinvestigators.org/downloads/), GWAS-EBI catalog (https://www.ebi.ac.uk/gwas/). GWAS summary statistics for PDAC were provided by the Pancreatic Cancer Cohort Consortium and the Pancreatic Cancer Case-Control Consortium.

#### GARFIELD analysis

We used GARFIELD algorithm ^65^ to calculate the enrichment of GWAS variants that overlapped with our cell type specific enhancer trees. Specifically, we used cell type-specific ATAC-seq peak coordinates that overlap with enhancer tree nodes (enhancers and promoters), and prepared annotation overlap files as required by the algorithm at different GWAS thresholds.

We used the data on LD tags, distance to TSS, and minor allele frequency provided by GARFIELD, which implements LD scores based on the UK10K project as the reference set for European population. GARFIELD enrichment tests were run individually using summary statistics from each of the GWAS studies listed above, and the coordinates for all four cell types (α, β, acinar, and duct) defined above were used as input annotations. The estimated odds ratios (ORs) and enrichment *p* values were computed at various GWAS *p* value thresholds (1 x 10^−05^,1 x 10^−06^,1 x 10^−07^, 5 x 10^−08^, and 1 x 10^−08^). The R code ‘**Garfield-Meff-Padj.R’** from GARFIELD was used to calculate an enrichment *p* value threshold. This threshold was adjusted using Bonferroni correction (*P* < 0.01) to account for multiple testing, based on the effective number of annotations (Meff = 4.45). The negative log2 of enrichment *p-*values for the tested GWAS summary statistics were plotted using ggplot package in R.

#### SNP-to-target identification and evaluating EPIC effect sizes

We used significantly enriched SNPs from the p < 1 x 10^−05^ GWAS threshold which was the largest bin containing the highest number of enriched SNPs from all four GWAS *p*-value thresholds. While this is a relatively lenient GWAS *p*-value threshold, we wanted to include the maximum number of enriched SNPs in the beginning for our gene mapping analysis. We then identified the overlapping SNPs by intersecting the SNP coordinates with enhancer tree node coordinates using BEDtools. Each overlap was represented with a unique identifier for each SNP-enhancer node-promoter link. The list was further filtered to include enhancer trees containing at least one enriched SNP node and has cell type-specific promoters that has an ESS > 0.7 ^47^. We focused on four trait-cell type pairings:

**T2D-β:** T2D SNP enriched β-cell trees
**PDAC-duct:** PDAC SNP enriched duct cell trees
**T2D-acinar:** T2D SNP enriched acinar cell trees
**PDAC-acinar:** PDAC SNP enriched acinar cell trees

We used EPIC to determine the effect size of all enhancer nodes that were included in these lists. Next, the EPIC evaluated nodes were stratified into two groups: SNP-overlapping or not-overlapping, and the effect sizes of these nodes were represented by group using CDF() function in R.

#### Visualization of select SNP-target gene regions

Examples of SNP-target gene networks for T2D and PDAC in acinar cells were visualized using locuszoom (https://my.locuszoom.org/) and the UCSC genome browser (https://genome.ucsc.edu/) on hg38 genome build.

## Author Contributions

L.W. and H.E.A. conceived and supervised the study; L.W., T.T., J.W., K.G. performed experiments; L.W, J.W., K.G, D.S., S.B., G.P., E.C., J.H., H.E.A performed computational analysis; J.H. and L.A. supervised GWAS enrichment analysis, S.B. generated the tree models and wrote the EPIC algorithm; L.W, S.B., H.E.A. analyzed and interpreted the findings; L.W., S.B and H.E.A. wrote the manuscript with inputs from all other co-authors.

## ACKNOWLEDGMENTS

We thank Y. Dalal, G. Hager, T. Misteli, S. Oberdoerffer, D. Larson, P. Batista, P. Rocha, J. Smith, W. Cui, and members of the Arda and Amundadottir laboratories for their help and critical review of this manuscript. We thank the Center for Cancer Research, National Cancer Institute core facilities— Genomics Core (Bethesda, MD), Sequencing Facility (Frederick, MD), Flow Cytometry Core (Bethesda, MD), and High-Throughput Imaging Facility (Bethesda, MD). This work used the computational resources of the NIH High Performance Cluster (Biowulf, https://hpc.nih.gov). This work was supported by the Intramural Research Program of the NIH, National Cancer Institute, Center for Cancer Research (grant no. ZIA BC011798 to H.E.A.). The American Cancer Society (ACS) funds the creation, maintenance, and updating of the Cancer Prevention Study II cohort. The authors express sincere appreciation to all Cancer Prevention Study-II participants, and to each member of the study and biospecimen management group. The authors would like to acknowledge the contribution to this study from central cancer registries supported through the Centers for Disease Control and Prevention’s National Program of Cancer Registries and cancer registries supported by the National Cancer Institute’s Surveillance Epidemiology and End Results Program. Where authors are identified as personnel of the International Agency for Research on Cancer / World Health Organization, the authors alone are responsible for the views expressed in this article and they do not necessarily represent the decisions, policy or views of the International Agency for Research on Cancer / World Health Organization. We acknowledge funding for the Women’s Health Study (WHS) source of data: CA047988, CA182913, HL043851, HL080467 and HL099355. We acknowledge WHI investigators listed here: https://www-whi-org.s3.us-west-2.amazonaws.com/wp-content/uploads/WHI-Investigator-Short-List.pdf. The WHI program is funded by the National Heart, Lung, and Blood Institute, National Institutes of Health, U.S. Department of Health and Human Services through 75N92021D00001, 75N92021D00002, 75N92021D00003, 75N92021D00004, 75N92021D00005. Human islets for research were provided by Integrated Islet Distribution Program (IIDP, NIH Grant # 2UC4DK098085) and the Alberta Diabetes Institute Islet Core at the University of Alberta in Edmonton (http://www.bcell.org/adi-isletcore.html) with the assistance of the Human Organ Procurement and Exchange (HOPE) program, Trillium Gift of Life Network (TGLN), and other Canadian organ procurement organizations. We gratefully acknowledge organ donors and their families.

## The Pancreatic Cancer Cohort Consortium

Jun Zhong^1^, Demetrius Albanes^2^, Gabriella Andreotti^3^, Alan A Arslan^4^, Laura Beane-Freeman^3^, Sonja I Berndt^3^, Julie E Buring^5,6^, Daniele Campa^7^, Federico Canzian^8^, Stephen J Chanock^9^, Yu Chen^10^, Sandra M Colorado-Yohar^11,12,13^, A. Heather Eliassen^14^, J. Michael Gaziano^15,16,17^, Graham G Giles^18,19,20^, Phyllis J Goodman^21^, Christopher A Haiman^22^, Mattias Johansson^23^, Verena Katzke^24^, Charles Kooperberg^25^, Peter Kraft^26^, Manolis Kogevinas^27,28,29,30^, I-Min Lee^5,6^, Loic LeMarchand^31^, Núria Malats^32,33^, Satu Männistö^34^, Marjorie L McCullough^35^, Roger Milne^18,19,20^, Stephen C Moore^2^, Lorelei Mucci^36^, Salvatore Panico^37^, Alpa V Patel^35^, Ulrike Peters^38^, Miquel Porta^29^, Francisco X Real^39,33,30^, Howard D Sesso^15,6^, Xiao-Ou Shu^40^, Meir J Stampfer^14,41^, Geoffrey S Tobias^9^, Kala Visvanathan^42,43^, Elisabete Weiderpass^44^, Nicolas Wentzensen^9^, Emily White^45,46^, Chen Yuan^47^, Wei Zheng^40^, Jean Wactawski-Wende^48^, Rachael Z Stolzenberg-Solomon^2^, Brian M Wolpin^49^, Laufey T Amundadottir^1^

^1^Laboratory of Translational Genomics, Division of Cancer Epidemiology and Genetics, National Cancer Institute, National Institutes of Health, Bethesda, MD, USA, ^2^Metabolic Epidemiology Branch, Division of Cancer Epidemiology and Genetics, National Cancer Institute, National Institutes of Health, Bethesda, MD, USA, ^3^Occupational and Environmental Epidemiology Branch, Division of Cancer Epidemiology and Genetics, National Cancer Institute, National Institutes of Health, Bethesda, MD, USA, ^4^Departments of Obstetrics and Gynecology and Population Health, NYU Grossman School of Medicine, NYU Perlmutter Comprehensive Cancer Center, New York, NY, USA, ^5^Division of Preventive Medicine, Department of Medicine, Brigham and Women’s Hospital, Boston, MA, USA, ^6^Department of Epidemiology, Harvard T.H. Chan School of Public Health, Boston, MA, USA, ^7^Unit of Genetics., Department of Biology, University of Pisa, Pisa, Italy, ^8^Genomic Epidemiology Group, German Cancer Research Center (DKFZ), Heidelberg, Germany, ^9^Division of Cancer Epidemiology and Genetics, National Cancer Institute, National Institutes of Health, Bethesda, MD, USA, ^10^Department of Population Health, NYU Grossman School of Medicine, NYU Perlmutter Comprehensive Cancer Center, New York, NY, USA, ^11^Department of Epidemiology, Murcia Regional Health Council, IMIB-Arrixaca, Murcia, Spain, ^12^CIBER Epidemiología y Salud Pública (CIBERESP), Spain, ^13^Research Group on Demography and Health, National Faculty of Public Health,, University of Antioquia, Medellín, Colombia, ^14^Department of Epidemiology, Harvard T. H. Chan School of Public Health, Boston, MA, USA, ^15^Division of Preventive Medicine, Brigham and Women’s Hospital, Boston, MA, USA, ^16^Division of Aging, Brigham and Women’s Hospital, Boston, MA, USA, ^17^Boston VA Healthcare System, Boston, MA, USA, ^18^Cancer Epidemiology Division, Cancer Council Victoria, Melbourne, VIC, Australia, ^19^Centre for Epidemiology and Biostatistics, Melbourne School of Population and Global Health, The University of Melbourne, Parkville, VIC, Australia, ^20^Precision Medicine, School of Clinical Sciences at Monash Health, Monash University, Melbourne, VIC, Australia, ^21^SWOG Statistical Center, Fred Hutchinson Cancer Research Center, Seattle, WA, USA, ^22^Department of Preventive Medicine, Keck School of Medicine, University of Southern California, Los Angeles, CA, ^23^Genomic Epidemiology Branch, International Agency for Research on Cancer (IARC/WHO), Lyon, France, ^24^Division of Cancer Epidemiology, German Cancer Research Center (DKFZ), Heidelberg, Germany, ^25^Division of Public Health Sciences, Fred Hutchinson Cancer Research Center, Seattle, WA, ^26^Trans-Divisional Research Program (TDRP), Division of Cancer Epidemiology and Genetics, National Cancer Institute, National Institutes of Health, Bethesda, MD, USA, ^27^ISGlobal, Centre for Research in Environmental Epidemiology (CREAL), Barcelona, Spain, ^28^CIBER Epidemiología y Salud Pública (CIBERESP), Barcelona, Spain, ^29^Hospital del Mar Institute of Medical Research (IMIM), Universitat Autònoma de Barcelona, Barcelona, Spain, ^30^Universitat Pompeu Fabra (UPF), Barcelona, Spain, ^31^Cancer Epidemiology Program, University of Hawaii Cancer Center, Honolulu, HI, USA, ^32^Genetic and Molecular Epidemiology Group, Spanish National Cancer Research Center (CNIO), Madrid, Spain, ^33^CIBERONC, Madrid, Spain, ^34^Department of Public Health and Welfare, Finnish Institute for Health and Welfare (THL), Helsinki, Finland, ^35^Department of Population Science, American Cancer Society, Atlanta, GA, USA, ^36^Department of Epidemiology, Harvard T. H. Chan School of Public Health, Boston, MA, ^37^Dipartimento Di Medicina Clinica E Chirurgia, Federico II University, Naples, Italy, ^38^Division of Public Health Sciences, Fred Hutchinson Cancer Center, Seattle, WA, USA, ^39^Epithelial Carcinogenesis Group, Molecular Oncology Programme, Spanish National Cancer Research Center (CNIO), Madrid, Spain, ^40^Division of Epidemiology, Department of Medicine, Vanderbilt Epidemiology Center, Vanderbilt-Ingram Cancer Center, Vanderbilt University School of Medicine, Nashville, TN, USA, ^41^Department of Nutrition, Harvard T. H. Chan School of Public Health, Boston, MA, USA, ^42^Department of Epidemiology, Johns Hopkins School of Public Health, Baltimore, MD, ^43^Department of Oncology, Sidney Kimmel Comprehensive Cancer Center, Johns Hopkins School of Medicine, Baltimore, MD, ^44^International Agency for Research on Cancer (IARC/WHO), Lyon, France, ^45^Division of Public Health Sciences, Fred Hutchinson Cancer Research Center, Seattle, WA, USA, ^46^Department of Epidemiology, University of Washington, Seattle, WA, USA, ^47^Department of Medical Oncology, Dana-Farber Cancer Institute, Harvard Medical School, Harvard University, Boston, MA, ^48^Department of Epidemiology and Environmental Health, University of Buffalo, Buffalo, NY, USA, ^49^Department of Medical Oncology, Dana-Farber Cancer Institute, Harvard Medical School, Harvard University, Boston, MA, USA

## Pancreatic Cancer Case-Control Consortium

Samuel O Antwi^1^, Paige M Bracci^2^, Steven Gallinger^3^, Michael Goggins^4^, Manal Hassan^5^, Elizabeth A Holly^2^, Rayjean J Hung^3^, Donghui Li^5^, Núria Malats^6,7^, Rachel E Neale^8^, Kari G Rabe^9^, Harvey A Risch^10^, Herbert Yu^11^, Alison P Klein^12,13,4^

^1^Department of Quantitative Health Sciences, Mayo Clinic College of Medicine, Jacksonville, FL, USA, ^2^Department of Epidemiology and Biostatistics, University of California, San Francisco, San Francisco, CA, USA, ^3^Lunenfeld-Tanenbaum Research Institute, Sinai Health System and University of Toronto, Toronto, Canada, ^4^Department of Pathology, Sol Goldman Pancreatic Cancer Research Center, Johns Hopkins School of Medicine, Baltimore, MD, USA, ^5^Department of Gastrointestinal Medical Oncology, University of Texas MD Anderson Cancer Center, Houston, TX, USA, ^6^Genetic and Molecular Epidemiology Group, Spanish National Cancer Research Center (CNIO), Madrid, Spain, ^7^CIBERONC, Madrid, Spain, ^8^Population Health Program, QIMR Berghofer Medical Research Institute, Brisbane, Australia, ^9^Department of Quantitative Health Sciences, Mayo Clinic College of Medicine, Rochester, MN, USA, ^10^Department of Chronic Disease Epidemiology, Yale School of Public Health, New Haven, CT, USA, ^11^Epidemiology Program, University of Hawaii Cancer Center, Honolulu, HI, USA, ^12^Department of Epidemiology, Johns Hopkins School of Public Health, Baltimore, MD, USA, ^13^Department of Oncology, Sidney Kimmel Comprehensive Cancer Center, Johns Hopkins School of Medicine, Baltimore, MD, USA

---

**Supplementary Figure 1 (relates to Figure 1):**
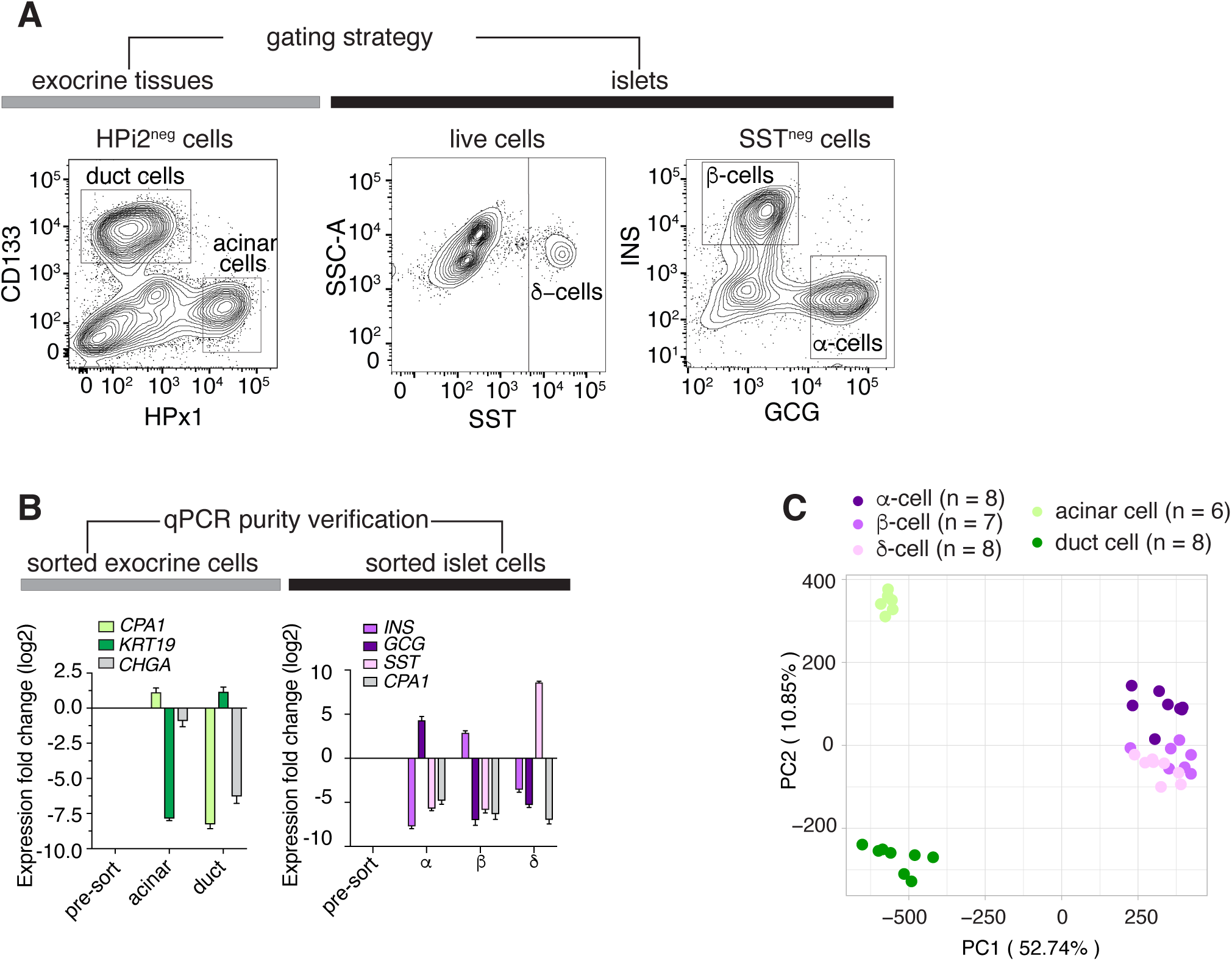
A. FACS plots showing the gating strategy to isolate different pancreas cell populations. Also see Supplementary Table 2. B. Bar plots showing the enrichment and depletion of marker genes in each purified cell population as determined by qPCR analysis. Results were normalized to pre-sorted cells. C. The Principal Component Analysis shows the clustering of ATAC-seq samples based on chromatin accessibility profiles across different pancreatic cell types. Each point represents a sample, and the samples are color-coded according to their respective cell type. The number of donors (n) used for each cell type is indicated.

**Supplementary Figure 2 (relates to Figure 2).**
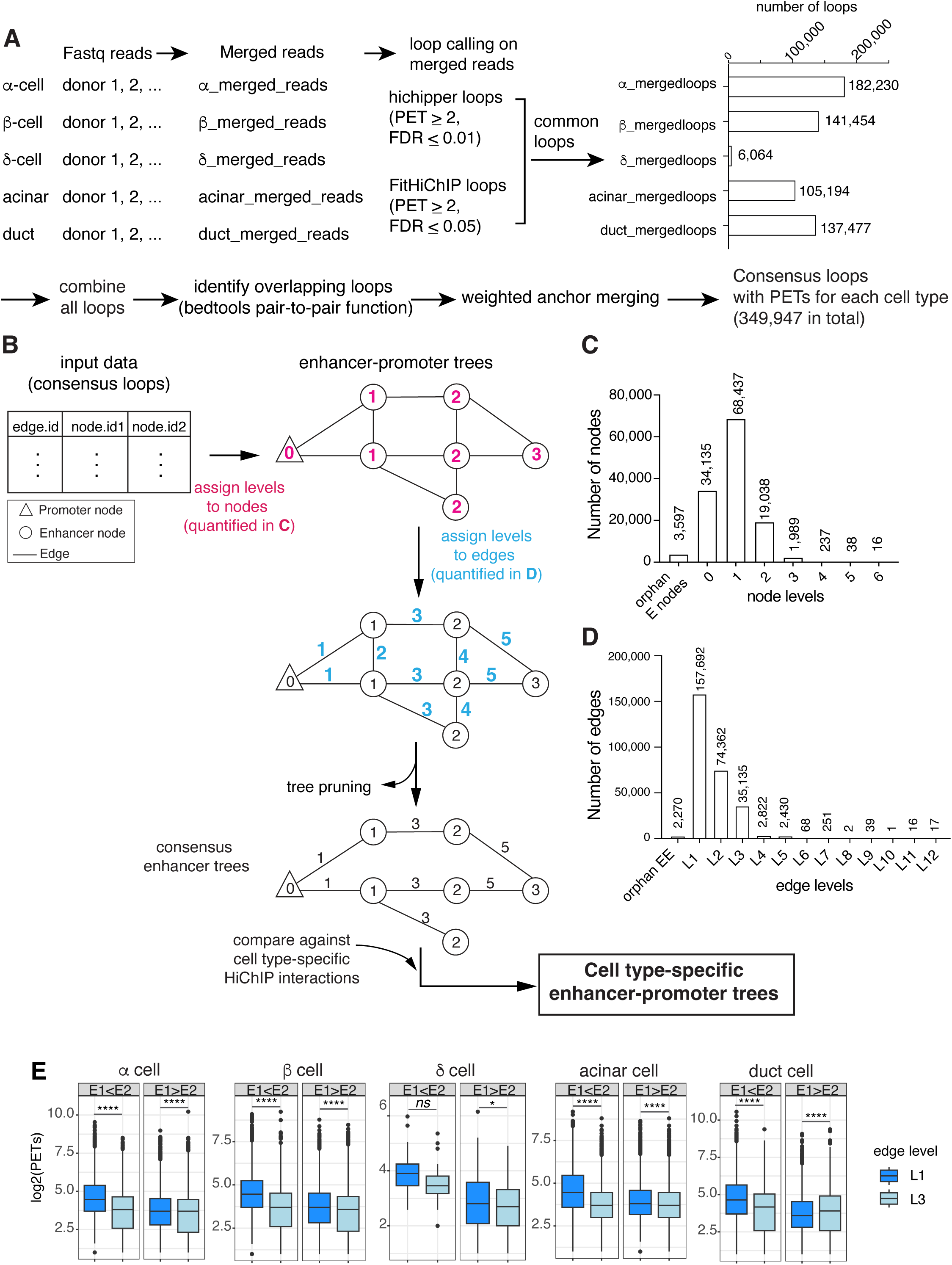
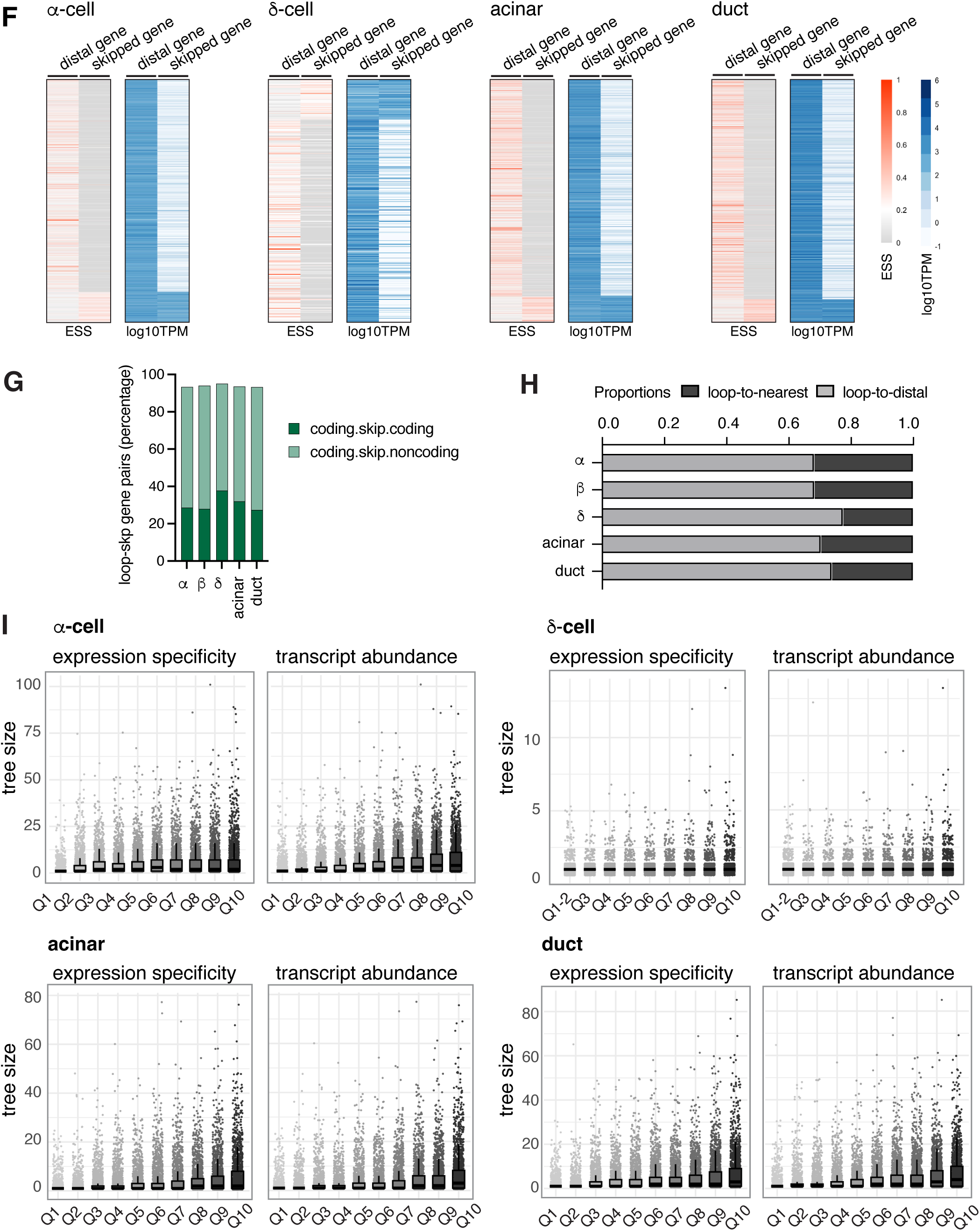
A. Flowchart describing consensus loop development. B. Schematic detaling enhancer tree construction based on consensus loops. C-D. Bar graphs depict the distribution of nodes (C) and edges (D) by connectivity level in enhancer trees before pruning. E. Interaction frequency stratified by distance of E1 and E2 relative to their corresponding promoters. Mann-Whitney test, **** *P*-value < 0.0001; * *P*-value < 0.05; ns, not significant. F. Heat maps show the expression specificity (ESS, red) and abundance (blue) of distally looped or skipped genes. Each row represents a gene pair that are either distally looped to or skipped by the same enhancer in α-, δ-, acinar or duct cells. G. Distribution of gene pairs based on their type in distally looping and skipped gene interactions across different pancreatic cell types (α-, β-, δ-, acinar or duct cells.). The dark green bars represent the percentage of gene pairs where both the distally looping and skipped genes are coding genes (coding.skip.coding). The light green bars show the percentage of pairs where the distally looping gene is a coding gene, but the skipped gene is a non-coding gene (coding.skip.noncoding). H. Fraction of enhancers looping to the nearest gene (dark grey) or a distal gene (light grey) in each cell type when noncoding genes are excluded from the datasets. I. Box plots depict the relationship between transcript abundance and the size of enhancer-promoter trees as measured by the number of enhancers linked to a single promoter. The x-axis represents the quantiles of expression specificity (ESS) or transcript abundance. The individual data points represent specific tree sizes for genes within each quantile.

**Supplementary Figure 3 (relates to Figure 3).**
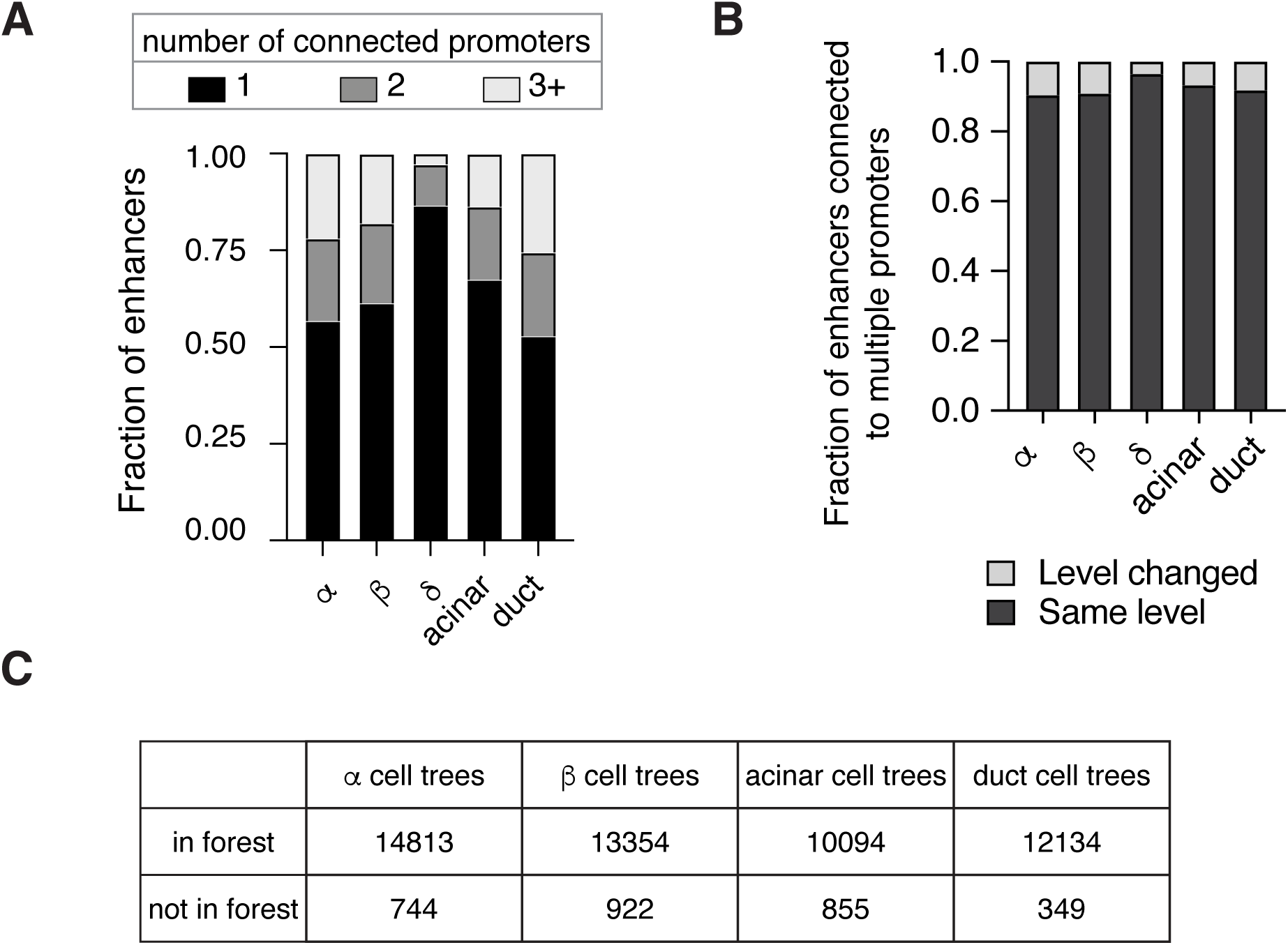
A. Bar graph shows fraction of enhancers connecting to one, two, three or more promoters quantified in each pancreatic cell type. B. Bar graph shows the proportion of enhancers that connect to multiple promoters in terms of level changes in each cell type. C. Table showing the number of trees belonging to a forest, stratified by cell type.

**Supplementary Figure 4 (relates to Figure 5).**
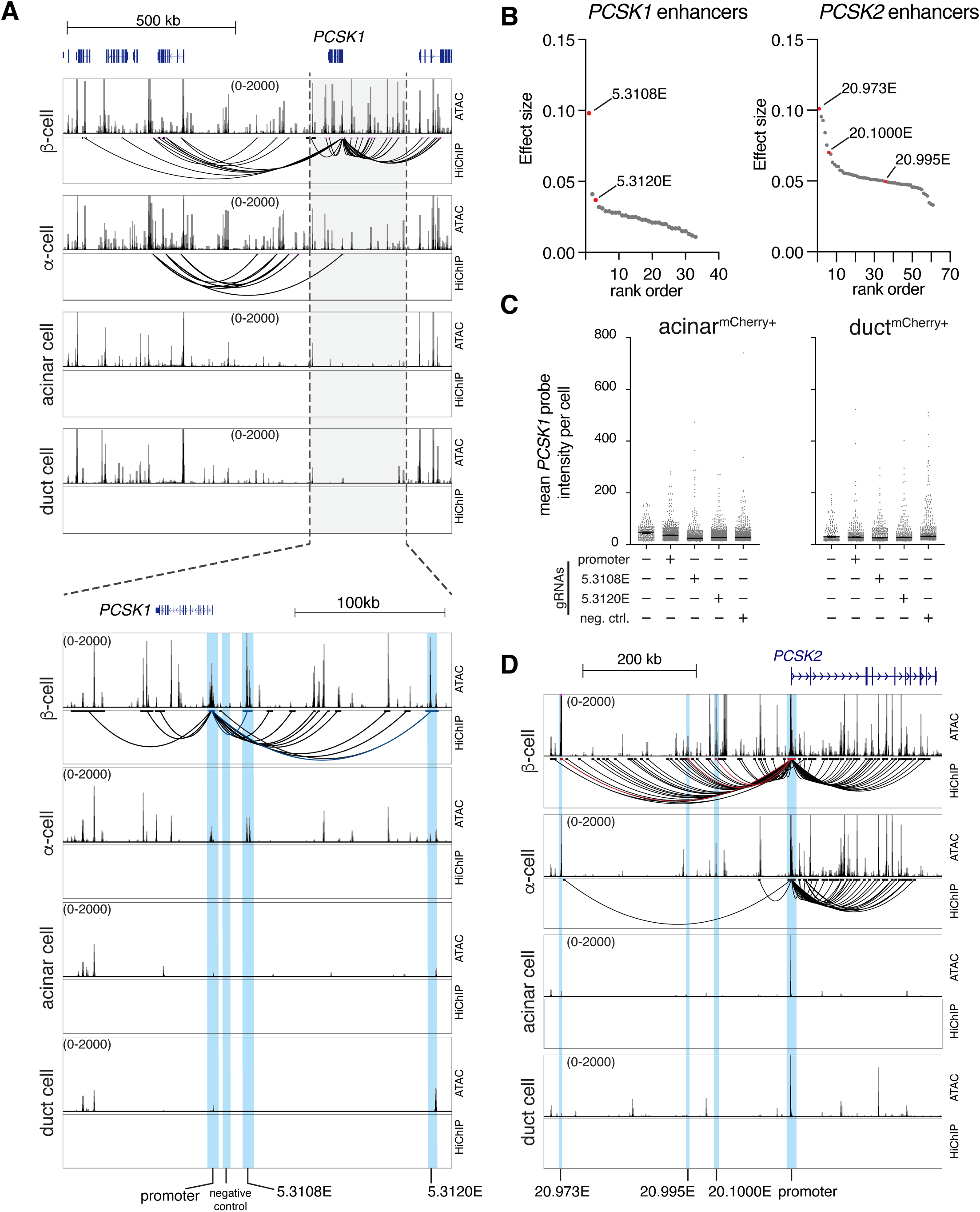
A. UCSC genome browser tracks displaying ATAC-seq peaks and HiChIP loops at the *PCSK1* locus across different pancreas cell types. The top set of tracks shows a broader region around the *PCSK1* gene, showing all the enhancer-promoter interactions corresponding to the *PCSK1* tree in β-cells. The bottom set of tracks zooms in on a region around *PCSK1,* blue highlighted regions indicate the top-ranking enhancers 5.3108E and 5.3120E, the promoter and the negative control regions. B. Scatter plots displaying the ranked effect sizes of β-cell *PCSK1* tree enhancers (left) and α-cell *PCSK2* tree enhancers (right). Each point represents an enhancer, ordered by effect size. The enhancers that were tested in CRISPR perturbation assays are marked in red. C. Quantification of *PCSK1* transcript levels in mCherry^+^ acinar cells and mCherry^+^ duct cells after CRISPRa targeting. Mean *PCSK1* probe intensity per cell is shown for cells treated with gRNAs targeting the promoter (positive control), enhancers 5.3108E and 5.3120E, and a negative control region. No activation was observed in either cell type or perturbation condition. n(mCherry^+^ acinar cells)= 4714, n(mCherry^+^ duct cells)= 2193, results were reproduced by at least two independent donors. D. UCSC genome browser tracks displaying ATAC-seq peaks and HiChIP loops at the *PCSK2* locus across different pancreas cell types. Blue highlighted regions indicate the promoter and EPIC-prioritized enhancers regions.

**Supplementary Figure 5 (relates to Figure 5).**
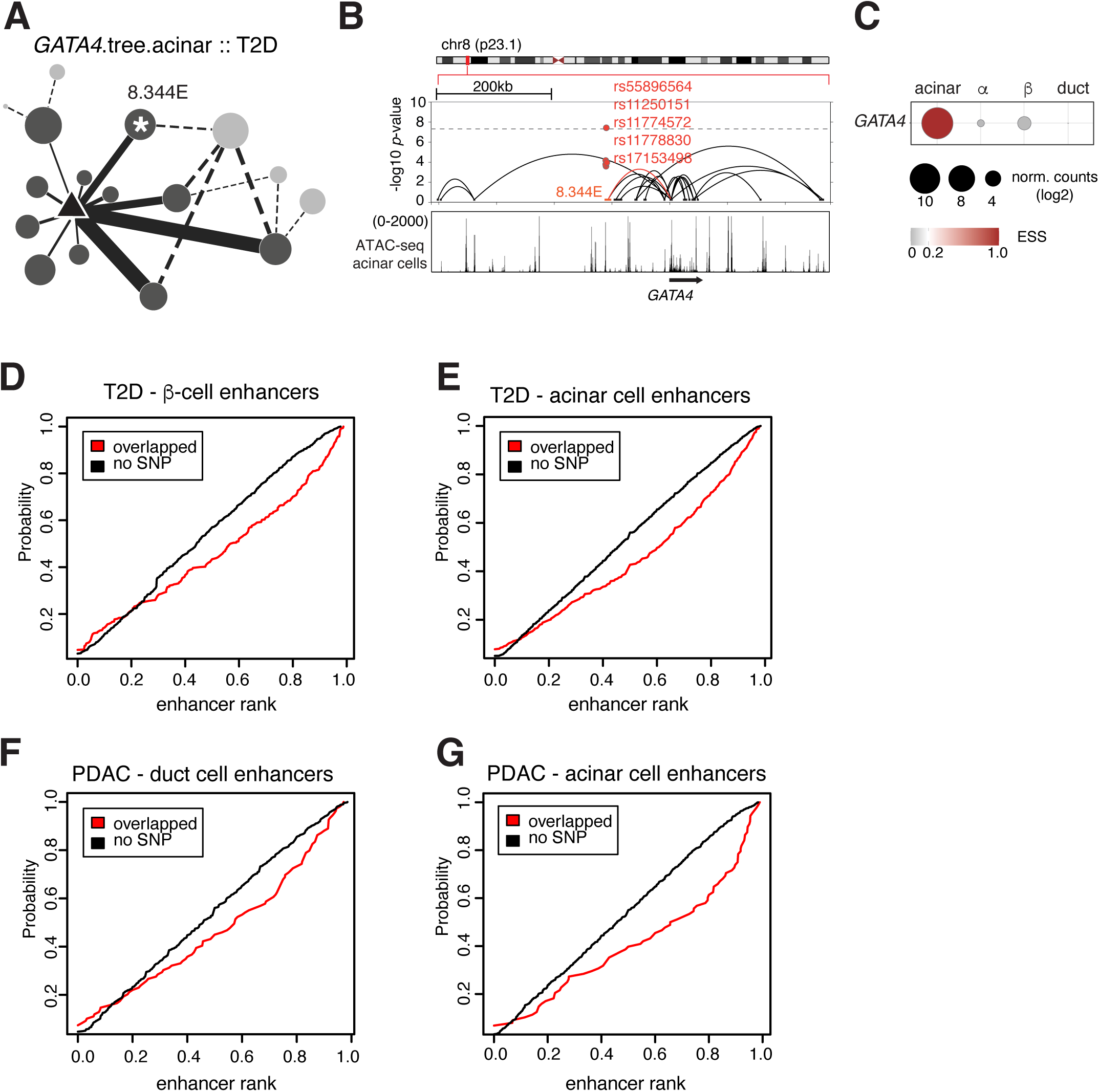
A. Network representation of the *GATA4* enhancer tree in acinar cells. Nodes represent enhancers, with the size of each node reflecting the ATAC-seq tag density and the thickness of the lines (edges) indicating the strength of the enhancer-promoter interactions as detected by HiChIP. The node with the asterisk indicates the SNP enriched enhancer. B. Combined UCSC genome browser and locus zoom plots displaying the enhancer tree elements at the *GATA4* locus in acinar cells. The locus zoom plot highlighting significant SNPs in red. The UCSC genome browser tracks below show the corresponding ATAC-seq peaks and HiChIP loops detected in acinar cells. Red highlights the node enriched with the significant SNPs. C. Bubble plot depicts the expression specificity (ESS) and transcript abundance (normalized counts) of *GATA4* transcripts in human pancreas cells based on single-cell RNA-seq data. D-G. Cumulative Distribution Function (CDF) plots comparing the enhancer ranks based on EPIC’s prioritization, with and without SNP overlap, for different cell types and GWAS traits. The top-ranking enhancer has the value of 1, and bottom-ranking enhancer has the value of zero. (D) The red line represents enhancers that overlap with SNPs associated with type 2 diabetes (T2D) in β-cells, while the black line represents enhancers without SNP overlap. (E) The same comparison for enhancers in acinar cells associated with T2D. (F) Enhancers overlapping with SNPs (red) and those without SNP overlap (black) in ductal cells associated with pancreatic ductal adenocarcinoma (PDAC). (G) The same comparison for enhancers in acinar cells associated with PDAC.

## REFERENCES

1. Xiao, A.Y., Tan, M.L.Y., Wu, L.M., Asrani, V.M., Windsor, J.A., Yadav, D., and Petrov, M.S. (2016). Global incidence and mortality of pancreatic diseases: a systematic review, meta-analysis, and meta-regression of population-based cohort studies. Lancet Gastroenterol. Hepatol. 1, 45–55. 10.1016/S2468-1253(16)30004-8.

2. Khalaf, N., El-Serag, H.B., Abrams, H.R., and Thrift, A.P. (2021). Burden of Pancreatic Cancer: From Epidemiology to Practice. Clin. Gastroenterol. Hepatol. Off. Clin. Pract. J. Am. Gastroenterol. Assoc. 19, 876–884. 10.1016/j.cgh.2020.02.054.

3. GBD 2021 Diabetes Collaborators (2023). Global, regional, and national burden of diabetes from 1990 to 2021, with projections of prevalence to 2050: a systematic analysis for the Global Burden of Disease Study 2021. Lancet Lond. Engl. 402, 203–234. 10.1016/S0140-6736(23)01301-6.

4. Jennings, R.E., Berry, A.A., Strutt, J.P., Gerrard, D.T., and Hanley, N.A. (2015). Human pancreas development. Dev. Camb. Engl. 142, 3126–3137. 10.1242/dev.120063.

5. Ma, Z., Zhang, X., Zhong, W., Yi, H., Chen, X., Zhao, Y., Ma, Y., Song, E., and Xu, T. (2023). Deciphering early human pancreas development at the single-cell level. Nat. Commun. 14, 5354. 10.1038/s41467-023-40893-8.

6. Habener, J.F., and Stanojevic, V. (2012). α-cell role in β-cell generation and regeneration. Islets 4, 188–198. 10.4161/isl.20500.

7. Puri, S., Folias, A.E., and Hebrok, M. (2015). Plasticity and dedifferentiation within the pancreas: development, homeostasis, and disease. Cell Stem Cell 16, 18–31. 10.1016/j.stem.2014.11.001.

8. Yu, X.-X., and Xu, C.-R. (2020). Understanding generation and regeneration of pancreatic β cells from a single-cell perspective. Dev. Camb. Engl. 147, dev179051. 10.1242/dev.179051.

9. Yuan, S., Norgard, R.J., and Stanger, B.Z. (2019). Cellular Plasticity in Cancer. Cancer Discov. 9, 837–851. 10.1158/2159-8290.CD-19-0015.

10. Marstrand-Daucé, L., Lorenzo, D., Chassac, A., Nicole, P., Couvelard, A., and Haumaitre, C. (2023). Acinar-to-Ductal Metaplasia (ADM): On the Road to Pancreatic Intraepithelial Neoplasia (PanIN) and Pancreatic Cancer. Int. J. Mol. Sci. 24, 9946. 10.3390/ijms24129946.

11. Crawford, H.C., Scoggins, C.R., Washington, M.K., Matrisian, L.M., and Leach, S.D. (2002). Matrix metalloproteinase-7 is expressed by pancreatic cancer precursors and regulates acinar-to-ductal metaplasia in exocrine pancreas. J. Clin. Invest. 109, 1437–1444. 10.1172/JCI15051.

12. Wang, L., Xie, D., and Wei, D. (2019). Pancreatic Acinar-to-Ductal Metaplasia and Pancreatic Cancer. Methods Mol. Biol. Clifton NJ 1882, 299–308. 10.1007/978-1-4939-8879-2_26.

13. Espinet, E., Gu, Z., Imbusch, C.D., Giese, N.A., Büscher, M., Safavi, M., Weisenburger, S., Klein, C., Vogel, V., Falcone, M., et al. (2021). Aggressive PDACs Show Hypomethylation of Repetitive Elements and the Execution of an Intrinsic IFN Program Linked to a Ductal Cell of Origin. Cancer Discov. 11, 638–659. 10.1158/2159-8290.CD-20-1202.

14. Gopalan, V., Singh, A., Rashidi Mehrabadi, F., Wang, L., Ruppin, E., Arda, H.E., and Hannenhalli, S. (2021). A Transcriptionally Distinct Subpopulation of Healthy Acinar Cells Exhibit Features of Pancreatic Progenitors and PDAC. Cancer Res. 81, 3958–3970. 10.1158/0008-5472.CAN-21-0427.

15. Stanger, B.Z., and Hebrok, M. (2013). Control of cell identity in pancreas development and regeneration. Gastroenterology 144, 1170–1179. 10.1053/j.gastro.2013.01.074.

16. Arda, H.E., Benitez, C.M., and Kim, S.K. (2013). Gene regulatory networks governing pancreas development. Dev. Cell 25, 5–13. 10.1016/j.devcel.2013.03.016.

17. Bastidas-Ponce, A., Scheibner, K., Lickert, H., and Bakhti, M. (2017). Cellular and molecular mechanisms coordinating pancreas development. Dev. Camb. Engl. 144, 2873–2888. 10.1242/dev.140756.

18. Haigis, K.M., Cichowski, K., and Elledge, S.J. (2019). Tissue-specificity in cancer: The rule, not the exception. Science 363, 1150–1151. 10.1126/science.aaw3472.

19. Smith, E., and Shilatifard, A. (2014). Enhancer biology and enhanceropathies. Nat. Struct. Mol. Biol. 21, 210–219. 10.1038/nsmb.2784.

20. Cebola, I. (2019). Pancreatic Islet Transcriptional Enhancers and Diabetes. Curr. Diab. Rep. 19, 145. 10.1007/s11892-019-1230-6.

21. Pachano, T., Haro, E., and Rada-Iglesias, A. (2022). Enhancer-gene specificity in development and disease. Dev. Camb. Engl. 149, dev186536. 10.1242/dev.186536.

22. Watanabe, K., Stringer, S., Frei, O., Umićević Mirkov, M., de Leeuw, C., Polderman, T.J.C., van der Sluis, S., Andreassen, O.A., Neale, B.M., and Posthuma, D. (2019). A global overview of pleiotropy and genetic architecture in complex traits. Nat. Genet. 51, 1339–1348. 10.1038/s41588-019-0481-0.

23. Zaugg, J.B., Sahlén, P., Andersson, R., Alberich-Jorda, M., de Laat, W., Deplancke, B., Ferrer, J., Mandrup, S., Natoli, G., Plewczynski, D., et al. (2022). Current challenges in understanding the role of enhancers in disease. Nat. Struct. Mol. Biol. 29, 1148–1158. 10.1038/s41594-022-00896-3.

24. Schoenfelder, S., and Fraser, P. (2019). Long-range enhancer-promoter contacts in gene expression control. Nat. Rev. Genet. 20, 437–455. 10.1038/s41576-019-0128-0.

25. Stitzel, M.L., Sethupathy, P., Pearson, D.S., Chines, P.S., Song, L., Erdos, M.R., Welch, R., Parker, S.C.J., Boyle, A.P., Scott, L.J., et al. (2010). Global epigenomic analysis of primary human pancreatic islets provides insights into type 2 diabetes susceptibility loci. Cell Metab. 12, 443–455. 10.1016/j.cmet.2010.09.012.

26. Parker, S.C.J., Stitzel, M.L., Taylor, D.L., Orozco, J.M., Erdos, M.R., Akiyama, J.A., van Bueren, K.L., Chines, P.S., Narisu, N., NISC Comparative Sequencing Program, et al. (2013). Chromatin stretch enhancer states drive cell-specific gene regulation and harbor human disease risk variants. Proc. Natl. Acad. Sci. U. S. A. 110, 17921–17926. 10.1073/pnas.1317023110.

27. Bramswig, N.C., Everett, L.J., Schug, J., Dorrell, C., Liu, C., Luo, Y., Streeter, P.R., Naji, A., Grompe, M., and Kaestner, K.H. (2013). Epigenomic plasticity enables human pancreatic α to β cell reprogramming. J. Clin. Invest. 123, 1275–1284. 10.1172/JCI66514.

28. Pasquali, L., Gaulton, K.J., Rodríguez-Seguí, S.A., Mularoni, L., Miguel-Escalada, I., Akerman, İ., Tena, J.J., Morán, I., Gómez-Marín, C., van de Bunt, M., et al. (2014). Pancreatic islet enhancer clusters enriched in type 2 diabetes risk-associated variants. Nat. Genet. 46, 136–143. 10.1038/ng.2870.

29. Arda, H.E., Tsai, J., Rosli, Y.R., Giresi, P., Bottino, R., Greenleaf, W.J., Chang, H.Y., and Kim, S.K. (2018). A Chromatin Basis for Cell Lineage and Disease Risk in the Human Pancreas. Cell Syst. 7, 310–322.e4. 10.1016/j.cels.2018.07.007.

30. Varshney, A., Scott, L.J., Welch, R.P., Erdos, M.R., Chines, P.S., Narisu, N., Albanus, R.D., Orchard, P., Wolford, B.N., Kursawe, R., et al. (2017). Genetic regulatory signatures underlying islet gene expression and type 2 diabetes. Proc. Natl. Acad. Sci. U. S. A. 114, 2301–2306. 10.1073/pnas.1621192114.

31. Rai, V., Quang, D.X., Erdos, M.R., Cusanovich, D.A., Daza, R.M., Narisu, N., Zou, L.S., Didion, J.P., Guan, Y., Shendure, J., et al. (2020). Single-cell ATAC-Seq in human pancreatic islets and deep learning upscaling of rare cells reveals cell-specific type 2 diabetes regulatory signatures. Mol. Metab. 32, 109–121. 10.1016/j.molmet.2019.12.006.

32. Chiou, J., Geusz, R.J., Okino, M.-L., Han, J.Y., Miller, M., Melton, R., Beebe, E., Benaglio, P., Huang, S., Korgaonkar, K., et al. (2021). Interpreting type 1 diabetes risk with genetics and single-cell epigenomics. Nature 594, 398–402. 10.1038/s41586-021-03552-w.

33. Chiou, J., Zeng, C., Cheng, Z., Han, J.Y., Schlichting, M., Miller, M., Mendez, R., Huang, S., Wang, J., Sui, Y., et al. (2021). Single-cell chromatin accessibility identifies pancreatic islet cell type- and state-specific regulatory programs of diabetes risk. Nat. Genet. 53, 455–466. 10.1038/s41588-021-00823-0.

34. Schmitt, A.D., Hu, M., Jung, I., Xu, Z., Qiu, Y., Tan, C.L., Li, Y., Lin, S., Lin, Y., Barr, C.L., et al. (2016). A Compendium of Chromatin Contact Maps Reveals Spatially Active Regions in the Human Genome. Cell Rep. 17, 2042–2059. 10.1016/j.celrep.2016.10.061.

35. Greenwald, W.W., Chiou, J., Yan, J., Qiu, Y., Dai, N., Wang, A., Nariai, N., Aylward, A., Han, J.Y., Kadakia, N., et al. (2019). Pancreatic islet chromatin accessibility and conformation reveals distal enhancer networks of type 2 diabetes risk. Nat. Commun. 10, 2078. 10.1038/s41467-019-09975-4.

36. Miguel-Escalada, I., Bonàs-Guarch, S., Cebola, I., Ponsa-Cobas, J., Mendieta-Esteban, J., Atla, G., Javierre, B.M., Rolando, D.M.Y., Farabella, I., Morgan, C.C., et al. (2019). Human pancreatic islet three-dimensional chromatin architecture provides insights into the genetics of type 2 diabetes. Nat. Genet. 51, 1137–1148. 10.1038/s41588-019-0457-0.

37. Su, C., Gao, L., May, C.L., Pippin, J.A., Boehm, K., Lee, M., Liu, C., Pahl, M.C., Golson, M.L., Naji, A., et al. (2022). 3D chromatin maps of the human pancreas reveal lineage-specific regulatory architecture of T2D risk. Cell Metab. 34, 1394–1409.e4. 10.1016/j.cmet.2022.08.014.

38. Weng, C., Gu, A., Zhang, S., Lu, L., Ke, L., Gao, P., Liu, X., Wang, Y., Hu, P., Plummer, D., et al. (2023). Single cell multiomic analysis reveals diabetes-associated β-cell heterogeneity driven by HNF1A. Nat. Commun. 14, 5400. 10.1038/s41467-023-41228-3.

39. Arda, H.E., Li, L., Tsai, J., Torre, E.A., Rosli, Y., Peiris, H., Spitale, R.C., Dai, C., Gu, X., Qu, K., et al. (2016). Age-Dependent Pancreatic Gene Regulation Reveals Mechanisms Governing Human β Cell Function. Cell Metab. 23, 909–920. 10.1016/j.cmet.2016.04.002.

40. Blodgett, D.M., Nowosielska, A., Afik, S., Pechhold, S., Cura, A.J., Kennedy, N.J., Kim, S., Kucukural, A., Davis, R.J., Kent, S.C., et al. (2015). Novel Observations From Next-Generation RNA Sequencing of Highly Purified Human Adult and Fetal Islet Cell Subsets. Diabetes 64, 3172–3181. 10.2337/db15-0039.

41. Creyghton, M.P., Cheng, A.W., Welstead, G.G., Kooistra, T., Carey, B.W., Steine, E.J., Hanna, J., Lodato, M.A., Frampton, G.M., Sharp, P.A., et al. (2010). Histone H3K27ac separates active from poised enhancers and predicts developmental state. Proc. Natl. Acad. Sci. U. S. A. 107, 21931–21936. 10.1073/pnas.1016071107.

42. Andersson, R., and Sandelin, A. (2020). Determinants of enhancer and promoter activities of regulatory elements. Nat. Rev. Genet. 21, 71–87. 10.1038/s41576-019-0173-8.

43. Field, A., and Adelman, K. (2020). Evaluating Enhancer Function and Transcription. Annu. Rev. Biochem. 89, 213–234. 10.1146/annurev-biochem-011420-095916.

44. Ray-Jones, H., and Spivakov, M. (2021). Transcriptional enhancers and their communication with gene promoters. Cell. Mol. Life Sci. CMLS 78, 6453–6485. 10.1007/s00018-021-03903-w.

45. Preissl, S., Gaulton, K.J., and Ren, B. (2023). Characterizing cis-regulatory elements using single-cell epigenomics. Nat. Rev. Genet. 24, 21–43. 10.1038/s41576-022-00509-1.

46. Pancaldi, V. (2023). Network models of chromatin structure. Curr. Opin. Genet. Dev. 80, 102051. 10.1016/j.gde.2023.102051.

47. Sturgill, D., Wang, L., and Arda, H.E. (2024). PancrESS - a meta-analysis resource for understanding cell-type specific expression in the human pancreas. BMC Genomics 25, 76. 10.1186/s12864-024-09964-y.

48. Kempfer, R., and Pombo, A. (2020). Methods for mapping 3D chromosome architecture. Nat. Rev. Genet. 21, 207–226. 10.1038/s41576-019-0195-2.

49. McArthur, E., and Capra, J.A. (2021). Topologically associating domain boundaries that are stable across diverse cell types are evolutionarily constrained and enriched for heritability. Am. J. Hum. Genet. 108, 269–283. 10.1016/j.ajhg.2021.01.001.

50. Choi, H.M.T., Beck, V.A., and Pierce, N.A. (2014). Next-generation in situ hybridization chain reaction: higher gain, lower cost, greater durability. ACS Nano 8, 4284–4294. 10.1021/nn405717p.

51. Choi, H.M.T., Schwarzkopf, M., Fornace, M.E., Acharya, A., Artavanis, G., Stegmaier, J., Cunha, A., and Pierce, N.A. (2018). Third-generation in situ hybridization chain reaction: multiplexed, quantitative, sensitive, versatile, robust. Dev. Camb. Engl. 145, dev165753. 10.1242/dev.165753.

52. Larson, M.H., Gilbert, L.A., Wang, X., Lim, W.A., Weissman, J.S., and Qi, L.S. (2013). CRISPR interference (CRISPRi) for sequence-specific control of gene expression. Nat. Protoc. 8, 2180–2196. 10.1038/nprot.2013.132.

53. Gilbert, L.A., Horlbeck, M.A., Adamson, B., Villalta, J.E., Chen, Y., Whitehead, E.H., Guimaraes, C., Panning, B., Ploegh, H.L., Bassik, M.C., et al. (2014). Genome-Scale CRISPR-Mediated Control of Gene Repression and Activation. Cell 159, 647–661. 10.1016/j.cell.2014.09.029.

54. Goodge, K.A., and Hutton, J.C. (2000). Translational regulation of proinsulin biosynthesis and proinsulin conversion in the pancreatic beta-cell. Semin. Cell Dev. Biol. 11, 235–242. 10.1006/scdb.2000.0172.

55. Orskov, C., Holst, J.J., Poulsen, S.S., and Kirkegaard, P. (1987). Pancreatic and intestinal processing of proglucagon in man. Diabetologia 30, 874–881. 10.1007/BF00274797.

56. Friis-Hansen, L., Lacourse, K.A., Samuelson, L.C., and Holst, J.J. (2001). Attenuated processing of proglucagon and glucagon-like peptide-1 in carboxypeptidase E-deficient mice. J. Endocrinol. 169, 595–602. 10.1677/joe.0.1690595.

57. Turpeinen, H., Ortutay, Z., and Pesu, M. (2013). Genetics of the first seven proprotein convertase enzymes in health and disease. Curr. Genomics 14, 453–467. 10.2174/1389202911314050010.

58. Stijnen, P., Ramos-Molina, B., O’Rahilly, S., and Creemers, J.W.M. (2016). PCSK1 Mutations and Human Endocrinopathies: From Obesity to Gastrointestinal Disorders. Endocr. Rev. 37, 347–371. 10.1210/er.2015-1117.

59. Smeekens, S.P., Montag, A.G., Thomas, G., Albiges-Rizo, C., Carroll, R., Benig, M., Phillips, L.A., Martin, S., Ohagi, S., and Gardner, P. (1992). Proinsulin processing by the subtilisin-related proprotein convertases furin, PC2, and PC3. Proc. Natl. Acad. Sci. U. S. A. 89, 8822–8826. 10.1073/pnas.89.18.8822.

60. Furuta, M., Carroll, R., Martin, S., Swift, H.H., Ravazzola, M., Orci, L., and Steiner, D.F. (1998). Incomplete processing of proinsulin to insulin accompanied by elevation of Des-31,32 proinsulin intermediates in islets of mice lacking active PC2. J. Biol. Chem. 273, 3431–3437. 10.1074/jbc.273.6.3431.

61. Yang, Y., Hua, Q.-X., Liu, J., Shimizu, E.H., Choquette, M.H., Mackin, R.B., and Weiss, M.A. (2010). Solution structure of proinsulin: connecting domain flexibility and prohormone processing. J. Biol. Chem. 285, 7847–7851. 10.1074/jbc.C109.084921.

62. Klein, A.P., Wolpin, B.M., Risch, H.A., Stolzenberg-Solomon, R.Z., Mocci, E., Zhang, M., Canzian, F., Childs, E.J., Hoskins, J.W., Jermusyk, A., et al. (2018). Genome-wide meta-analysis identifies five new susceptibility loci for pancreatic cancer. Nat. Commun. 9, 556. 10.1038/s41467-018-02942-5.

63. Chen, J., Spracklen, C.N., Marenne, G., Varshney, A., Corbin, L.J., Luan, J., Willems, S.M., Wu, Y., Zhang, X., Horikoshi, M., et al. (2021). The trans-ancestral genomic architecture of glycemic traits. Nat. Genet. 53, 840–860. 10.1038/s41588-021-00852-9.

64. Mahajan, A., Spracklen, C.N., Zhang, W., Ng, M.C.Y., Petty, L.E., Kitajima, H., Yu, G.Z., Rüeger, S., Speidel, L., Kim, Y.J., et al. (2022). Multi-ancestry genetic study of type 2 diabetes highlights the power of diverse populations for discovery and translation. Nat. Genet. 54, 560–572. 10.1038/s41588-022-01058-3.

65. Iotchkova, V., Ritchie, G.R.S., Geihs, M., Morganella, S., Min, J.L., Walter, K., Timpson, N.J., UK10K Consortium, Dunham, I., Birney, E., et al. (2019). GARFIELD classifies disease-relevant genomic features through integration of functional annotations with association signals. Nat. Genet. 51, 343–353. 10.1038/s41588-018-0322-6.

66. Shaw-Smith, C., De Franco, E., Lango Allen, H., Batlle, M., Flanagan, S.E., Borowiec, M., Taplin, C.E., van Alfen-van der Velden, J., Cruz-Rojo, J., Perez de Nanclares, G., et al. (2014). GATA4 mutations are a cause of neonatal and childhood-onset diabetes. Diabetes 63, 2888–2894. 10.2337/db14-0061.

67. Song, M., Pebworth, M.-P., Yang, X., Abnousi, A., Fan, C., Wen, J., Rosen, J.D., Choudhary, M.N.K., Cui, X., Jones, I.R., et al. (2020). Cell-type-specific 3D epigenomes in the developing human cortex. Nature 587, 644–649. 10.1038/s41586-020-2825-4.

68. Murphy, D., Salataj, E., Di Giammartino, D.C., Rodriguez-Hernaez, J., Kloetgen, A., Garg, V., Char, E., Uyehara, C.M., Ee, L.-S., Lee, U., et al. (2024). 3D Enhancer-promoter networks provide predictive features for gene expression and coregulation in early embryonic lineages. Nat. Struct. Mol. Biol. 31, 125–140. 10.1038/s41594-023-01130-4.

69. Bergman, D.T., Jones, T.R., Liu, V., Ray, J., Jagoda, E., Siraj, L., Kang, H.Y., Nasser, J., Kane, M., Rios, A., et al. (2022). Compatibility rules of human enhancer and promoter sequences. Nature 607, 176–184. 10.1038/s41586-022-04877-w.

70. Martinez-Ara, M., Comoglio, F., and van Steensel, B. (2023). Large-scale analysis of the integration of enhancer-enhancer signals by promoters. 10.7554/elife.91994.1.

71. van Mierlo, G., Pushkarev, O., Kribelbauer, J.F., and Deplancke, B. (2023). Chromatin modules and their implication in genomic organization and gene regulation. Trends Genet. TIG 39, 140–153. 10.1016/j.tig.2022.11.003.

72. Mian, Y., Wang, L., Keikhosravi, A., Guo, K., Misteli, T., Arda, H.E., and Finn, E.H. (2024). Cell type- and transcription-independent spatial proximity between enhancers and promoters. Mol. Biol. Cell 35, ar96. 10.1091/mbc.E24-02-0082.

73. Luppino, J.M., and Joyce, E.F. (2020). Single cell analysis pushes the boundaries of TAD formation and function. Curr. Opin. Genet. Dev. 61, 25–31. 10.1016/j.gde.2020.03.005.

74. Dekker, J., Alber, F., Aufmkolk, S., Beliveau, B.J., Bruneau, B.G., Belmont, A.S., Bintu, L., Boettiger, A., Calandrelli, R., Disteche, C.M., et al. (2023). Spatial and temporal organization of the genome: Current state and future aims of the 4D nucleome project. Mol. Cell 83, 2624–2640. 10.1016/j.molcel.2023.06.018.

75. Antal, C.E., Oh, T.G., Aigner, S., Luo, E.-C., Yee, B.A., Campos, T., Tiriac, H., Rothamel, K.L., Cheng, Z., Jiao, H., et al. (2023). A super-enhancer-regulated RNA-binding protein cascade drives pancreatic cancer. Nat. Commun. 14, 5195. 10.1038/s41467-023-40798-6.

76. Bevacqua, R.J., Zhao, W., Merheb, E., Kim, S.H., Marson, A., Gloyn, A.L., and Kim, S.K. (2024). Multiplexed CRISPR gene editing in primary human islet cells with Cas9 ribonucleoprotein. iScience 27, 108693. 10.1016/j.isci.2023.108693.

77. Wolpin, B.M., Rizzato, C., Kraft, P., Kooperberg, C., Petersen, G.M., Wang, Z., Arslan, A.A., Beane-Freeman, L., Bracci, P.M., Buring, J., et al. (2014). Genome-wide association study identifies multiple susceptibility loci for pancreatic cancer. Nat. Genet. 46, 994–1000. 10.1038/ng.3052.

78. López de Maturana, E., Rodríguez, J.A., Alonso, L., Lao, O., Molina-Montes, E., Martín-Antoniano, I.A., Gómez-Rubio, P., Lawlor, R., Carrato, A., Hidalgo, M., et al. (2021). A multilayered post-GWAS assessment on genetic susceptibility to pancreatic cancer. Genome Med. 13, 15. 10.1186/s13073-020-00816-4.

79. Chiou, J., Geusz, R.J., Okino, M.-L., Han, J.Y., Miller, M., Melton, R., Beebe, E., Benaglio, P., Huang, S., Korgaonkar, K., et al. (2021). Interpreting type 1 diabetes risk with genetics and single-cell epigenomics. Nature 594, 398–402. 10.1038/s41586-021-03552-w.

80. Chandra, V., Bhattacharyya, S., Schmiedel, B.J., Madrigal, A., Gonzalez-Colin, C., Fotsing, S., Crinklaw, A., Seumois, G., Mohammadi, P., Kronenberg, M., et al. (2021). Promoter-interacting expression quantitative trait loci are enriched for functional genetic variants. Nat. Genet. 53, 110–119. 10.1038/s41588-020-00745-3.

81. Corces, M.R., Trevino, A.E., Hamilton, E.G., Greenside, P.G., Sinnott-Armstrong, N.A., Vesuna, S., Satpathy, A.T., Rubin, A.J., Montine, K.S., Wu, B., et al. (2017). An improved ATAC-seq protocol reduces background and enables interrogation of frozen tissues. Nat. Methods 14, 959–962. 10.1038/nmeth.4396.

82. Buenrostro, J.D., Wu, B., Chang, H.Y., and Greenleaf, W.J. (2015). ATAC-seq: A Method for Assaying Chromatin Accessibility Genome-Wide. Curr. Protoc. Mol. Biol. 109, 21.29.1–21.29.9. 10.1002/0471142727.mb2129s109.

83. Smith, J.P., Corces, M.R., Xu, J., Reuter, V.P., Chang, H.Y., and Sheffield, N.C. (2021). PEPATAC: an optimized pipeline for ATAC-seq data analysis with serial alignments. NAR Genomics Bioinforma. 3, lqab101. 10.1093/nargab/lqab101.

84. Quinlan, A.R., and Hall, I.M. (2010). BEDTools: a flexible suite of utilities for comparing genomic features. Bioinforma. Oxf. Engl. 26, 841–842. 10.1093/bioinformatics/btq033.

85. Heinz, S., Benner, C., Spann, N., Bertolino, E., Lin, Y.C., Laslo, P., Cheng, J.X., Murre, C., Singh, H., and Glass, C.K. (2010). Simple combinations of lineage-determining transcription factors prime cis-regulatory elements required for macrophage and B cell identities. Mol. Cell 38, 576–589. 10.1016/j.molcel.2010.05.004.

86. Anders, S., and Huber, W. (2010). Differential expression analysis for sequence count data. Genome Biol. 11, R106. 10.1186/gb-2010-11-10-r106.

87. Mumbach, M.R., Rubin, A.J., Flynn, R.A., Dai, C., Khavari, P.A., Greenleaf, W.J., and Chang, H.Y. (2016). HiChIP: efficient and sensitive analysis of protein-directed genome architecture. Nat. Methods 13, 919–922. 10.1038/nmeth.3999.

88. Servant, N., Varoquaux, N., Lajoie, B.R., Viara, E., Chen, C.-J., Vert, J.-P., Heard, E., Dekker, J., and Barillot, E. (2015). HiC-Pro: an optimized and flexible pipeline for Hi-C data processing. Genome Biol. 16, 259. 10.1186/s13059-015-0831-x.

89. Lareau, C.A., and Aryee, M.J. (2018). hichipper: a preprocessing pipeline for calling DNA loops from HiChIP data. Nat. Methods 15, 155–156. 10.1038/nmeth.4583.

90. Bhattacharyya, S., Chandra, V., Vijayanand, P., and Ay, F. (2019). Identification of significant chromatin contacts from HiChIP data by FitHiChIP. Nat. Commun. 10, 4221. 10.1038/s41467-019-11950-y.

91. Doench, J.G., Hartenian, E., Graham, D.B., Tothova, Z., Hegde, M., Smith, I., Sullender, M., Ebert, B.L., Xavier, R.J., and Root, D.E. (2014). Rational design of highly active sgRNAs for CRISPR-Cas9-mediated gene inactivation. Nat. Biotechnol. 32, 1262–1267. 10.1038/nbt.3026.

92. Doench, J.G., Fusi, N., Sullender, M., Hegde, M., Vaimberg, E.W., Donovan, K.F., Smith, I., Tothova, Z., Wilen, C., Orchard, R., et al. (2016). Optimized sgRNA design to maximize activity and minimize off-target effects of CRISPR-Cas9. Nat. Biotechnol. 34, 184–191. 10.1038/nbt.3437.

93. Hsu, P.D., Scott, D.A., Weinstein, J.A., Ran, F.A., Konermann, S., Agarwala, V., Li, Y., Fine, E.J., Wu, X., Shalem, O., et al. (2013). DNA targeting specificity of RNA-guided Cas9 nucleases. Nat. Biotechnol. 31, 827–832. 10.1038/nbt.2647.

94. Gu, B., Swigut, T., Spencley, A., Bauer, M.R., Chung, M., Meyer, T., and Wysocka, J. (2018). Transcription-coupled changes in nuclear mobility of mammalian cis-regulatory elements. Science 359, 1050–1055. 10.1126/science.aao3136.

95. Jiang, L., Zheng, Z., Fang, H., and Yang, J. (2021). A generalized linear mixed model association tool for biobank-scale data. Nat. Genet. 53, 1616–1621. 10.1038/s41588-021-00954-4.

96. Schwartzentruber, J., Cooper, S., Liu, J.Z., Barrio-Hernandez, I., Bello, E., Kumasaka, N., Young, A.M.H., Franklin, R.J.M., Johnson, T., Estrada, K., et al. (2021). Genome-wide meta-analysis, fine-mapping and integrative prioritization implicate new Alzheimer’s disease risk genes. Nat. Genet. 53, 392–402. 10.1038/s41588-020-00776-w.

